# Spatial genomics maps the structure, character and evolution of cancer clones

**DOI:** 10.1101/2021.04.16.439912

**Authors:** Artem Lomakin, Jessica Svedlund, Carina Strell, Milana Gataric, Artem Shmatko, Jun Sung Park, Young Seok Ju, Stefan Dentro, Vitalii Kleshchevnikov, Vasyl Vaskivskyi, Tong Li, Omer Ali Bayraktar, Luiza Moore, Sarah Pinder, Andrea L Richardson, Peter J Campbell, Moritz Gerstung, Mats Nilsson, Lucy R Yates

**Author notes:** These authors contributed equally (Artem Lomakin, Jessica Svedlund).

## Abstract

Subclonality is a universal feature of cancers yet how clones grow, are spatially organised, differ phenotypically or influence clinical outcome is unclear. To address this, we developed base specific in situ sequencing (BaSISS). In fixed tissues, transcripts harbouring clone-defining mutations are detected, converted into quantitative clone maps and characterised through multi-layered data integration. Applied to 8 samples from key stages of breast cancer progression BaSISS localised 1.42 million genotype informative transcripts across 4.9cm^2^ of tissue. Microscopic clonal topographies are shaped by resident tissue architectures. Distinct transcriptional, histological and immunological features distinguish coexistent genetic clones. Spatial lineage tracing temporally orders clone features associated with the emergence of aggressive clinical traits. These results highlight the pivotal role of spatial genomics in deciphering the mechanisms underlying cancer progression.

## Introductions

Cancer growth is the result of mutation and selection of ever more proliferative clones analogous to Darwinian evolutionary theory(*1*–*3*). A consequence of this relentless process is that cancers are patchworks of genetically related but distinct groups of cells termed subclones(*4, 5*). While the somatic evolution model is well established due to the almost omnipresent existence of cancer subclones in bulk or multi-regional sequencing data(*4*–*8*), relatively little is currently known about the nature or causes of spatial patterns of cancer growth, phenotypic characteristics of distinct subclonal lineages or their interactions with the microenvironment. Still this information appears key because adverse cancer outcomes – growth, progression and recurrence – are properties of genetically distinct subclones (*7, 9*–*11*).

While a range of spatial molecular profiling strategies based on spatial RNA barcoding(*12, 13*), or fluorescence microscopy of single RNA molecules using different types of molecular probes exist(*14*–*17*), they do not perform the critical function of isolating genetic subclones in tissue context because gene expression profiles are highly plastic. Evolutionary cancer genomics has demonstrated that lineage tracing using somatic mutations is a powerful and highly specific tool for tracing the subclonal origins of aggressive disease in earlier lesions(*7, 18, 19*). Histology driven sampling, such as laser capture microdissection(*20*) combined with low input nucleic acid library sequencing or even single cell sequencing goes some way towards resolving subclone spatial structure(*21, 22*). However, even the most exhaustive sampling strategy will struggle to provide an unbiased representation of the cancer landscape particularly across larger areas. Recently it has been demonstrated that molecular probes can be targeted to detect RNA in a sequence specific fashion, enabling detection of individual mutant transcripts *in situ(15, 23)*. Still, these technologies have not been able to comprehensively map multiple clones, define spatial cancer evolution or subclone specific phenotypes.

To overcome these limitations, we have developed a Base Specific In Situ Sequencing (BaSISS) methodology that extends the In Situ Sequencing (ISS) protocol by incorporating multiplexed detection of clone specific mutations in fixed tissue specimens (*24, 25*). A dedicated Bayesian model then allows the interpretation of multi-layered spatial data in genetic clone-specific context. By applying the method to eight tissues from two multifocal breast cancers we generate the first ever large scale quantitative maps of cancer clones and three key messages emerge. 1) Patterns of spatial genetic heterogeneity are profoundly influenced by resident tissue structures; 2) Coexistent genetic clones can have distinct transcriptional, histological and immunological characteristics; 3) In preinvasive, invasive and locally metastatic breast tumours, the emergence of aggressive disease features can be temporally ordered and localised in genetic and histological contexts providing insights into the biology underlying cancer progression.

## Results

### BaSISS detects bespoke panels of cancer-specific mutations in fixed tissues

Cancer evolution produces multiple genetically related yet distinct clones each characterised by a unique combination of somatic mutations, known as the genotype, that are related by the underlying phylogenetic tree (*26*). The spatial patterns created by coexisting cancer clones have not previously been directly observed. We established a pipeline to allow spatial mapping of cancer clones detected through standard bulk whole genome sequencing (WGS) and mutation clustering approaches (**Fig. 1A, Supp. Fig. S1, Supp. Methods**)(*7, 9*). Representative mutations from each cluster/branch of the phylogenetic tree were selected for spatial detection using BaSISS. The approach uses multiplexed highly sequence-specific padlock oligonucleotide probes with target recognition arms and 4-5 nucleotide reader barcodes to detect both mutant and wild-type alleles of each target (**Table S1**). Padlock probes bind to complementary DNA (cDNA), undergo rolling circle amplification and are read using sequencing by ligation with fluorophore-labelled interrogation probes (*15, 27*–*29*).

**Fig. 1.**
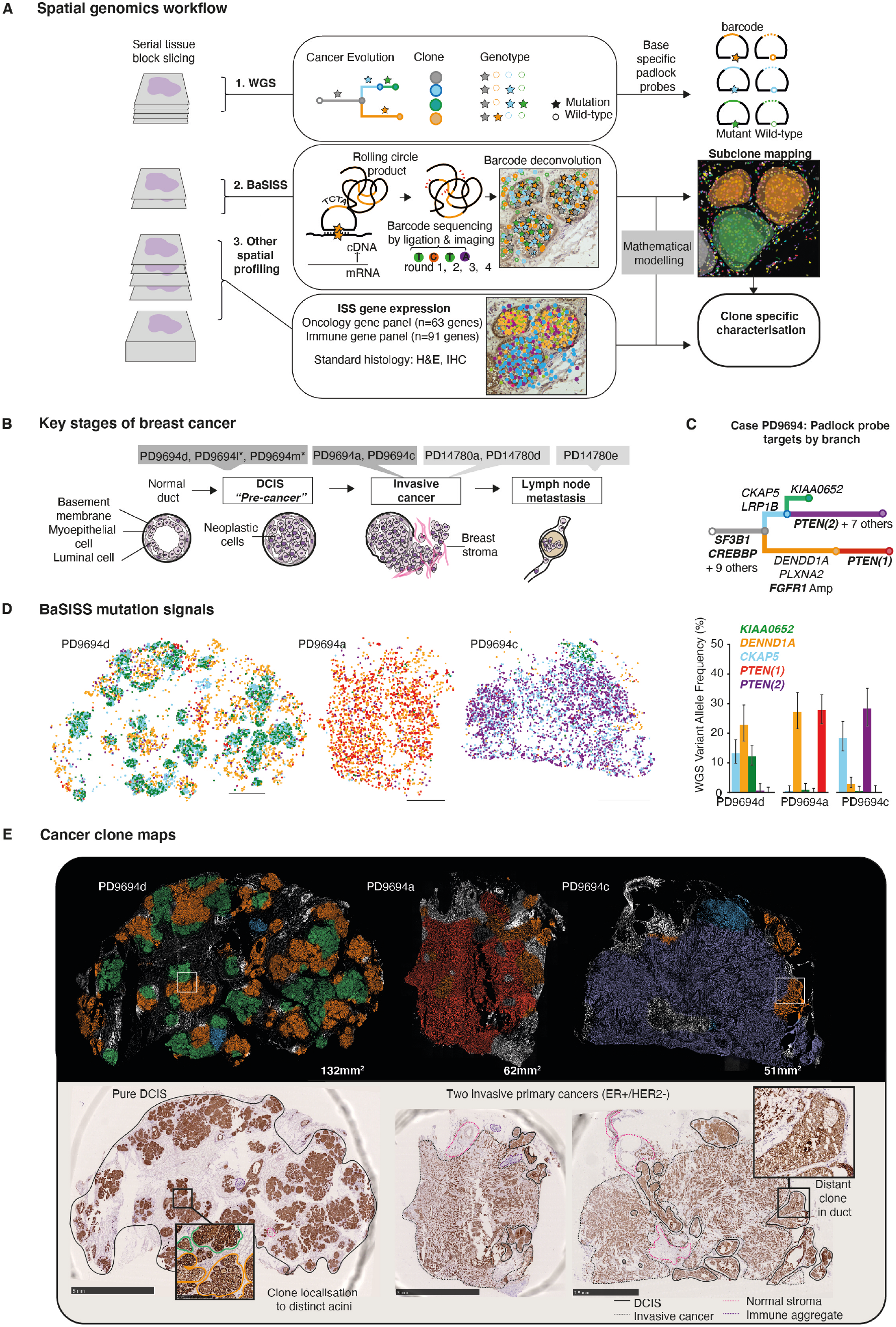
Spatial Genomics generates maps of cancer clones. (**A**) The Spatial Genomics workflow detects whole genome sequencing (WGS) defined subclonal mutations within serial tissue sections using base specific in-situ sequencing (BaSISS). For mathematical aspects see Fig. S1. (**B**) To develop the spatial genomics approach we selected two cases where multiple samples (PD*, shaded boxes) chart the distinct stages of primary breast cancer progression. Asterisk indicates no WGS data. (**C**) For case PD9694 most likely phylogenetic tree derived from WGS (samples PD9694a,c,d) and BaSISS data are presented with padlock probe targets named on each branch (driver mutations in bold). The barplot reveals the bulk genomic data (WGS plus targeted capture) derived variant allele frequency (VAF) of selected branch mutations in genes denoted in correspondingly coloured text (error bars report CIs = 5 - 95%). (**D**) BaSISS mutation signal plots – each dot represents the location of a mutation specific barcode, relating to the mutations reported in barplot (C). (**E**) Maps of the most prevalent clone projected on the DAPI image (reported if cancer cell fraction > 25% and inferred local cell density > 300 cells/mm2). Annotated pan-cytokeratin (pan-CK) IHC images of sequenced sections (bottom row, epithelial cells brown). Focus images show DCIS structure with coloured lines depicting approximate clone borders. DCIS = Ductal carcinoma in situ, ISS = gene expression in situ sequencing; IHC = immunohistochemistry, ER = oestrogen receptor, HER2 = Human epidermal growth factor receptor-2, cDNA = complementary DNA; mRNA = messenger RNA; H&E = haematoxylin and eosin.

To assess whether spatial mutation signals provide a meaningful representation of the bulk genomic data, we first applied it to three different regions of a breast tumour obtained from the same mastectomy (whole breast) specimen (case PD9694) with confirmed inter- and intra-sample genetic heterogeneity defined by WGS (**Fig. 1B-E**; **Table S2-3**)(*7*). We designed 51 BaSISS probes to target representative genetic variants from the phylogenetic tree trunk and branches (tree annotations; **Fig. 1C, Table S1**). BaSISS was applied to 10µm thick, fresh frozen tissue sections up to 1.75cm in diameter derived from the same tissue blocks used for WGS. On average, 96% of BaSISS reads were on target and the median number of signals reporting each target was 3,719 (combined mutant and wild-type) across the 3 samples (**Table S4, Supp. Methods**). Plotting BaSISS mutation signals coloured according to the relevant branch, reveals inter-sample differences and these are largely consistent with bulk genomic data derived variant allele frequencies (VAF) (R=0.48-0.61, Pearson’s) (**Fig. 1D, Fig. S2A-B)**. Replicating the BaSISS experiment on serial tissue samples generates highly concordant sample-wise VAFs (R=0.76-0.93, Pearson’s) and similar spatial signal distribution patterns (**Fig. S2C-D**). Spatial mutation and wild-type signal segregation patterns allowed the most likely phylogenetic tree, as represented in Figure 1C, to be selected from two solutions previously considered equally likely based solely on WGS data (**Fig. S2D-E, Supp. Methods**).

### Spatial genomics generates large scale cancer clone maps

To generate continuous spatial subclone maps that quantify the local composition of cancer subclones and normal cells we developed a statistical algorithm that exploits BaSISS signals as well as local cell counts (derived from the DAPI channel during the fluorescence microscopy of BaSISS) using two dimensional Gaussian processes (**Fig. S1**; **Supp. Methods**). The variational Bayesian model also accounts for unspecific or wrongly decoded BaSISS signals and variable probe efficiency and is augmented by VAFs in bulk genomic sequencing data from serial tissue sections. The resulting maps of cancer (and normal) clones were reconstructed across a scale of up to 132mm^2^ and with resolution of approximately 109µm (**Table S2**).

We applied the model to BaSISS data generated from eight tissue samples from 2 mastectomy specimens that together constitute 4.9cm^2^ of breast tissue (**Fig. 1B, Table S2**). The cases were selected to chart the landmark stages of breast cancer progression. The first case, PD9694, consists of two discrete oestrogen receptor (ER) positive invasive primary breast cancers with ductal carcinoma in situ (DCIS). DCIS is a non-obligate precursor of invasive breast cancer consisting of neoplastic cells restricted to the duct lumen. Breach through the basement membrane into the breast stroma marks progression to invasive cancer. The second case, PD14780, includes two ER negative invasive breast cancers and a draining axillary lymph node that contains metastatic cells.

Mapping the dominant clone across each tissue reveals striking genetic variation within and between samples from the same case (**Fig. 2-4**). Comparison with histological appearances confirms that the model correctly differentiates between neoplastic regions (coloured fields) and normal tissues (white DAPI nuclei revealed; **Fig. 1E**, frequency plots reach zero; **Fig. 2A**). Applying the model to the validation BaSISS data generates highly consistent results (**Fig. S3A**).

**Fig. 2.**
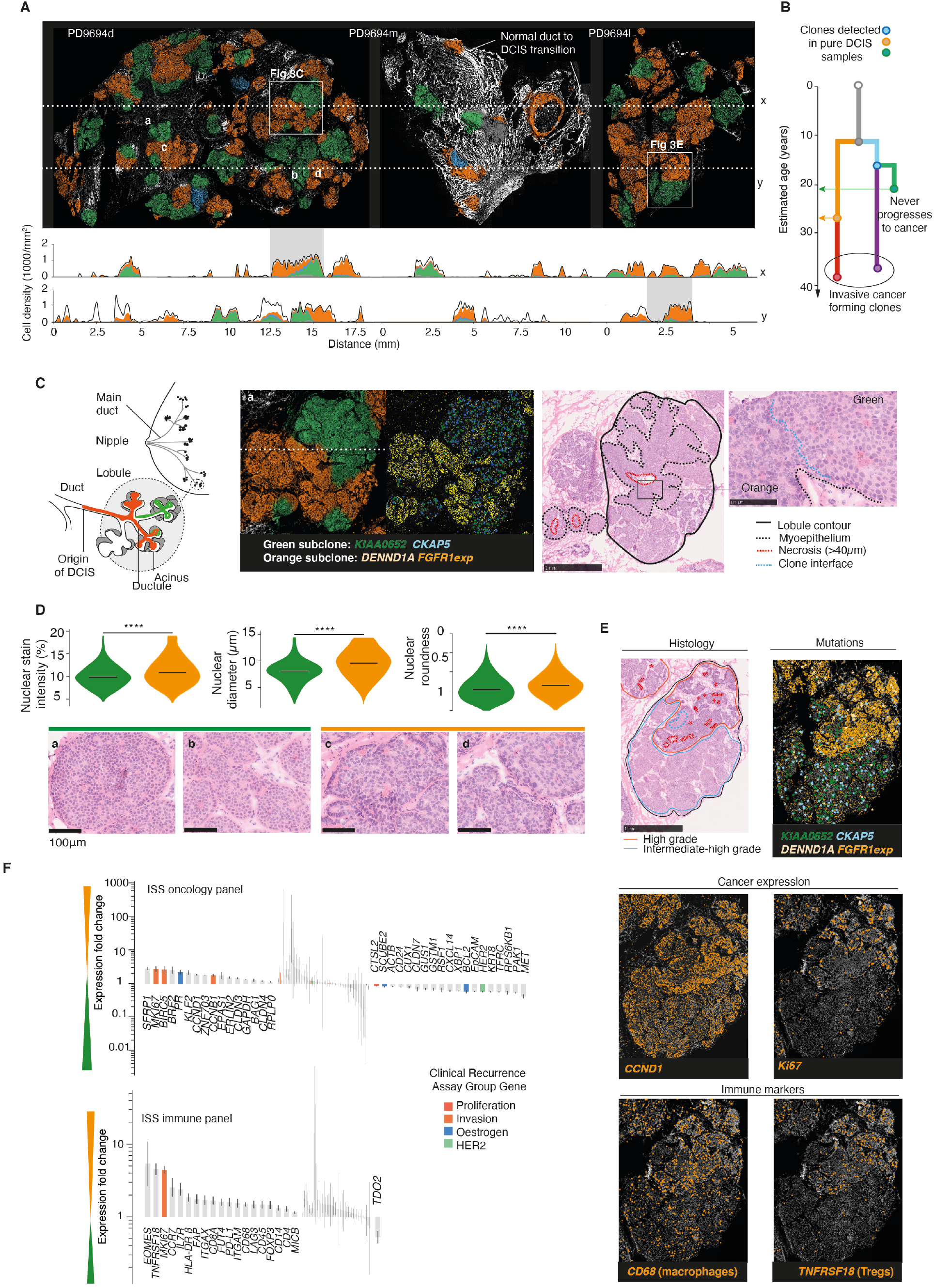
Histological, transcriptional and immunological differences between DCIS clones with different fates. (**A**) Maps of pure DCIS clones (as for Fig. 1E) and frequency plots of local, mean cancer (coloured areas) and non-cancer (white) composition, corresponding to horizontal dashed lines. (**B**) WGS derived phylogenetic tree reports relative timing of emergence (arrows) and evolutionary fates of DCIS clones in case PD9694. Timing estimates are derived from branch mutation burdens and patient age and assume steady mutation rates. (**C**) Pure DCIS clone growth patterns: Cartoon illustrates anatomical organisation of breast duct and lobule system and possible green/orange clone locations. Other images provide detailed views of segregated clone growth within a distended lobule in PD9694d (see A): clone fields (left) and select BaSISS mutation signals (middle) on DAPI image (coloured by clone: *FGFR1* is an expression probe so is more highly expressed but not exclusive to the orange subclone, other probes are mutation specific); annotated H&E stained serial section with focus image demonstrating different appearances across clone interface. (**D**) Histological appearances (H&E stained serial sections) of select areas corresponding to letters a-d in (A) reveal larger, more pleomorphic and intensely stained nuclei in orange clone. Violin plots report nuclear morphological features of green (11,365 nuclei) and orange (11,699 nuclei) regions (all differences significant, p < 0.0001, Mann-Whitney U). (**E**) BaSISS and ISS data projected on the DAPI image reveal orange clone specific characteristics in a selected area of PD9694l(A). (**F**) Barplots report fold changes in ISS gene panel expression signals between orange (1,515 tiles) and green clone regions (2,583 tiles) (predominant clone defines the location). Annotated genes have significant expression differences (probability of positive log-ratio (PPLR) after Bonferroni correction < 0.01). Genes are ordered by combined PPLR and direction of change (fine lines report CIs = 5 - 95%).

### Resident tissue structure influences subclonal growth patterns

Examining the samples from the three stages of cancer progression together, it is evident that the resident tissue structure plays a fundamental role in defining the observed patterns of intratumoral heterogeneity. One consequence is the intimate juxtaposition of genetically distant clones. For example, the lymph node sample (PD14780e) that is examined in detail in relation to Figure 4, contains multiple clones but the sinus spaces are monopolised by a single clone. Similarly, the purple, invasive primary cancer PD9694c, is studded by evolutionarily distant orange clones but these are intraductal populations of neoplastic cells that are physically separated from the main cancer mass by the duct membrane (inset box, PD9694c; **Fig. 1E**). Similarly, in the three ‘pure’ DCIS samples, the microscopic structure of ducts and lobules underlies the striking, seemingly random mosaic of green and orange clones that are predicted to have diverged decades before (inset box, PD9694d; **Fig. 1E, Fig. 2A-B)**.

The quantitative nature of these spatial genomics data allows us to further investigate the growth patterns in relation to tissue structure. In the pure DCIS samples (PD9694d,l,m), individual acinar and ductal ‘spaces’, defined by a myoepithelial cell layer and/or intervening stroma, are typically occupied by a complete clonal sweep (frequency plots; **Fig. 2A, Fig. S3B**). We deduce that this appearance is not simply a consequence of colonisation of distinct arms of the ductal system because we also observe distinct clonal fronts within several lobules that are corroborated by BaSISS mutation data, histological features and spatial transcriptomic data (**Fig. 2C-E**). These appearances might arise due to mutual tolerance or equal fitness, but in this case, as discussed below, the ability to spatially characterise co-existent clones provides evidence that we are more likely to be observing an incomplete clonal sweep by a fitter clone.

### Spatial genomics reveals characteristic differences between DCIS clones with different fates

Why some, but not all DCIS lesions progress to a potentially lethal invasive cancer is poorly understood. This question is usually addressed by comparing cohorts of DCIS samples with different clinical outcomes but this approach is inherently difficult due to the heterogeneity of breast cancer as a disease(*30, 31*). Here, we demonstrate that spatial genomics provides a novel approach to the problem, by allowing the comparison of related DCIS clones that share many genetic and all host features yet manifest different clinical outcomes. In case PD9694, an invasive cancer arose from the orange clone but the green clone, despite predating orange by several years, never progressed (**Fig. 2B**).

Consistent with a more aggressive phenotype the orange clone cells have larger, more pleomorphic and more intensely stained nuclei as confirmed by digital pathology (**Fig. 2C-D**). Clone-specific histological features are remarkably stable, being recapitulated within virtually every distinct space and are also appreciated at the single nucleus level where clones meet, adding weight to BaSISS model inferences (**Fig. 2C-D, Fig. S3C**). To derive clone-specific expression patterns, spatial transcriptomic data were generated on serial tissues sections using two bespoke ISS targeted panels: An immune marker panel (n=63 genes) and an oncology panel (n=91 genes) composed of a variety of breast epithelial markers and cancer pathway related genes including those derived from a clinical breast cancer recurrence assay (higher risk genes = proliferation, invasion, *HER2* group; lower risk genes = oestrogen group)(*32, 33*) (**Fig. 2F, Table S1, Table S5**). The per nucleus signal density was determined in regions dominated by either the orange or green subclone and significant differences determined (probability of positive log-ratio (PPLR) after Bonferroni correction < 0.01) (**Table S6, Supp. Methods**). Relative to green, the ill-fated orange clone expresses higher levels of cell cycle regulatory oncogenes *CCND1, CCNB1*, the cell survival factor *Survivin* and the proliferative marker *Ki67* (1.8-4.4 fold) (**Fig. 2E-F, Fig. S3D**). Proliferative markers have demonstrated prognostic and predictive value in various clinical studies of DCIS progression risk (*33*–*35*).

Given the barrier between DCIS and the surrounding stroma, we were surprised to also observe that the majority of immune panel signals are enriched within the bounds of the orange clone fields. The most significantly enriched immune marker genes include those associated with T-regulatory cells (*TNFRSF18, ITGAM*), macrophages (*CD68*) and immune checkpoint inhibition (*PD-L1, LAG3)* (1.5-4.4 fold). The fibroblast marker (*CD34*) and fibroblast activation protein, *FAP*, signals are also denser in these areas (**Fig. 2F, Fig. S3E**). These findings are consistent with various studies that propose that the stromal environment plays an active role in driving DCIS progression but provides new evidence that this mechanism is clone specific (*36*–*40*).

Raw ISS signals overlaid on the histological image provide visual confirmation of the highly clone-specific nature of some signals and furthermore, the within clone variation in spatial localisation (**Fig. 2E**). It is remarkable that many of the features that differentiate between precancerous clones with different evolutionary trajectories foreshadow, albeit at lower amplitude, many of the changes that distinguish invasive cancers from preinvasive cancers in general (**Fig. 3**) (*36, 41, 42*).

**Fig. 3.**
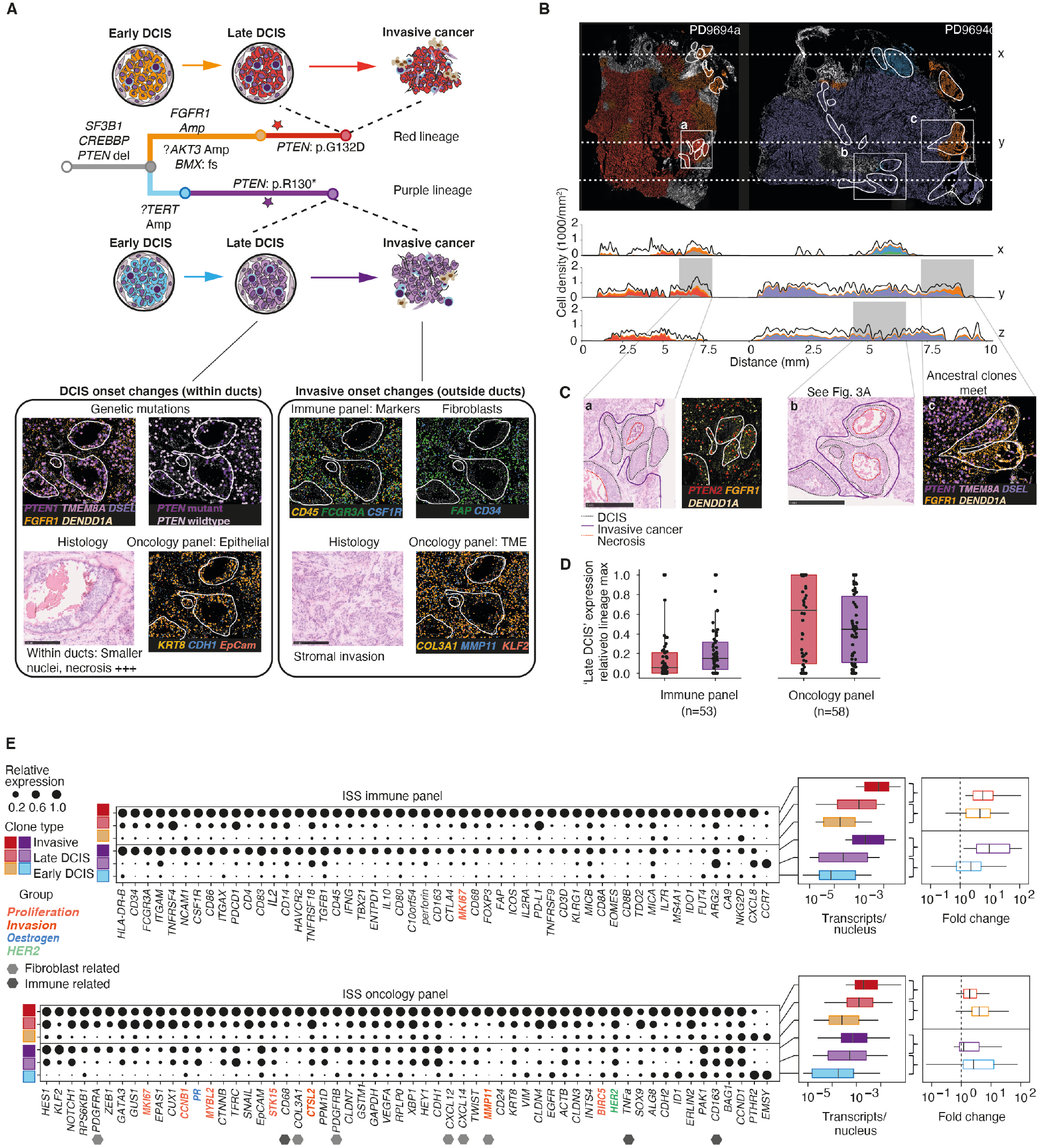
Temporal ordering of genetic, transcriptional and immunological changes during the development of invasive cancer. (**A**) Cartoon illustrates parallel lineages of progression from DCIS to invasive cancer. Potential driver mutations annotate relevant branches of the phylogenetic tree. DCIS onset changes (left box) include *PTEN* driver mutations and expression of genes typically associated with epithelial/cancer origin. Invasive onset changes (right box) include expression of genes associated with the tumour microenvironment (TME). (**B**) Clone maps (as per Fig. 1E) and frequency plots of invasive cancers with DCIS regions marked in white: *PTEN* mutant clones (red/purple) occupy several mm2 of histologically confirmed DCIS (A and **C**). (**D**) Boxplots report, for the genes upregulated between early DCIS and invasive cancer, the extent to which this is acquired within late DCIS. Significantly altered genes are defined as having PPLR after Bonferroni correction < 0.01. (**E**) A dot plot showing the relative expression of the same genes from (D) at the different progression stages in the 2 lineages. Dot area = (transcript/nucleus) divided by maximum value for each gene. Boxplots report transcripts/ nucleus (left) for genes shown and fold change between indicated comparisons (right). For all boxplots we report median, lower and upper quartiles (box) and 5-95 percentiles (whiskers). Expression per clone was defined on 136 - 3262 tiles, see Table S6 for details.

### Temporal ordering of genetic, transcriptional and immunological changes during the development of invasive cancer

How DCIS progresses to an invasive cancer is poorly understood. In the previous section we observed characteristic differences between competing clones in pure DCIS samples and here we extend this by tracing the changes accompanying the progression to invasive cancer – a potentially fatal condition (**Fig. 3A**). The two separate invasive cancers (PD9694a and PD9694c) arose from a common precursor clone along divergent lineages – orange to red and blue to purple – with parallel loss of *PTEN* through distinct damaging mutations (**Fig. 3A-B**). *PTEN* loss is the only invasion specific driver mutation identified within each lineage and was evidently under strong selection during progression as has been demonstrated in other breast cancers (*43*). In both lineages, the *PTEN* mutant clone dominates the invasive cancer but unexpectedly, also accounts for sizable areas of DCIS consistent with an intraductal onset (**Fig. 3B-C**). The terminal purple lineage clone is differentiated by 8 private mutations and these are all present at similar levels in the DCIS and invasive compartments supporting previous reports of genetic similarity between the two disease states (*21, 30*) (**Fig. S4A**). Histological appearances are more consistent with this reflecting genetic evolution within the ducts rather than widespread recolonisation by the invasive cancer and is referred to herein as ‘late DCIS’ to differentiate it from the ancestral DCIS clone (early DCIS). By comparing the targeted ISS gene expression data in early DCIS, late DCIS (with *PTEN* mutations), and invasive cancer clones along each lineage we sought to temporally order when the non-genetic features associated with an invasive cancer emerge (**Fig. 3A**). Here, we investigate to which extent transcriptional changes are replicated by the two lineages that display convergent genetic evolution in the same host and same genetic background.

**Fig. 4.**
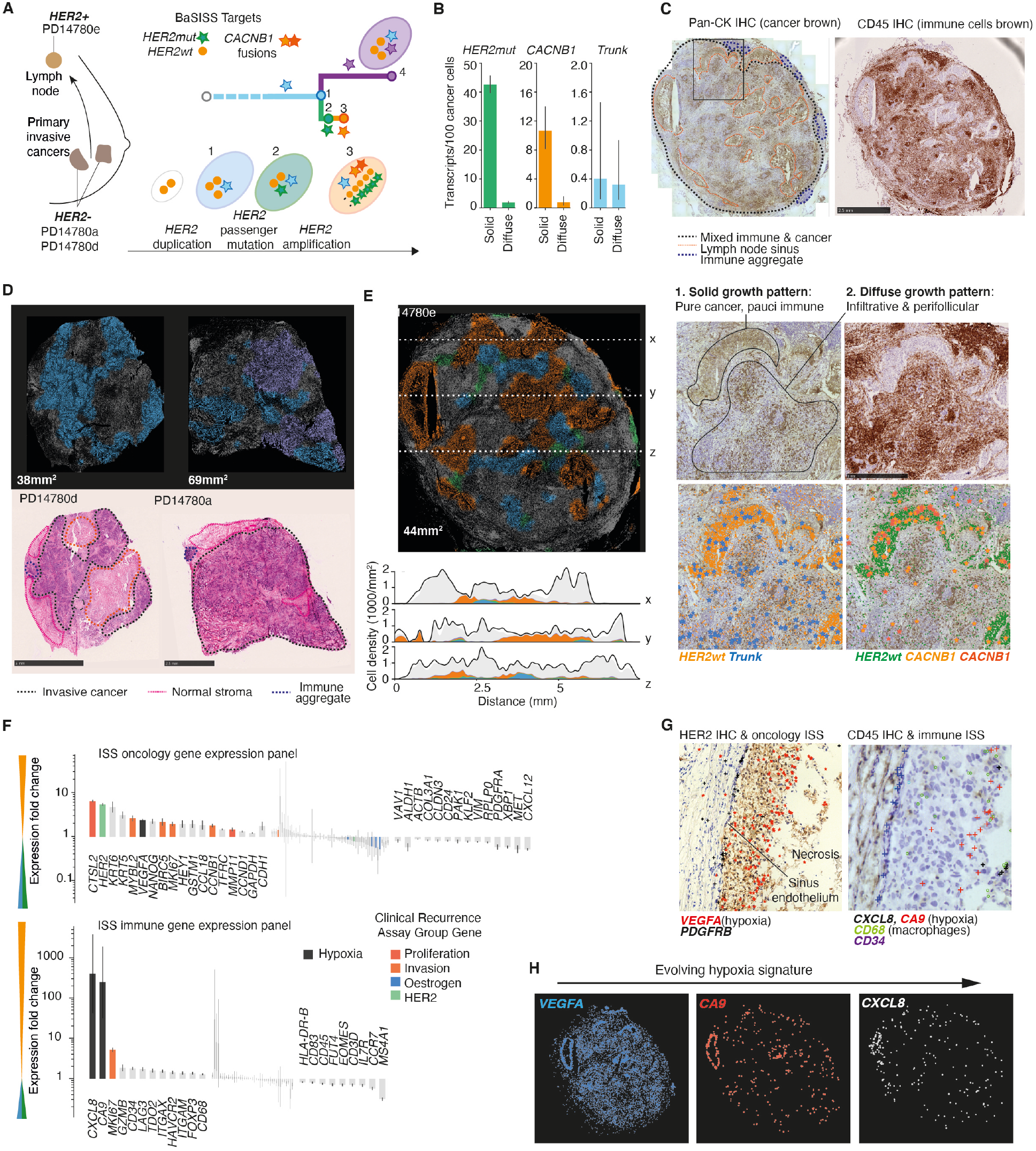
Tracing aggressive disease in a lymph node metastasis. (**A**) Case PD14780 samples, phylogenetic tree and genotypes (ovals represent a cell) inferred from WGS data and BaSISS signals. (**B**) Barplot of BaSISS signals extracted from 939 and 894 tiles in solid and diffuse growth pattern regions provide evidence of at least two genotypes: One with amplified *HER2* mutation signals and *CACNB1* fusion signals, the other with low level/ absent *HER2* mutation signals but similar trunk mutation signal density (CIs = 5-95%). ‘Trunk’ comprises 3 different mutations. (**C**) Sample PD14780e, BaSISS/ ISS sequenced lymph node sections stained by IHC with pan-cytokeratin (pan-CK, stains cancer cells brown) and CD45 (stains immune cells brown). Whole section (top row), focus area (middle row) and focus area (pan-CK stained) with overlaid BaSISS signals. (**D**-**E**) BaSISS subclone fields (top row) (DAPI projected, colour reflects the most prevalent subclone, reported if cancer cell fraction > 15% (D) and > 25% (E)) and (bottom row) annotated H&E stained serial sections (D) or subclone frequency plots (E). (**F**) Barplots of subclone specific gene expression fold changes in PD14780e (orange versus blue/green aggregated signals) for immune and oncology panels ordered by PPLR and direction of change. Genes with significant expression difference (PPLR after Bonferroni correction < 0.01) are annotated. (**G**) Spatial patterns of *PDGFRB, CD34* and hypoxia related ISS signals in PD14780e overlaid on HER2 (left) and CD45 IHC stained sections(right). (**H**) Spatial patterns of 3 hypoxia related genes show spatial patterns and are cumulative.

For each lineage, gene expression profiles were determined for the early DCIS, late DCIS and invasive cancer regions using targeted ISS. For 53 immune panel and 58 oncology panel genes, a significant difference in expression is evident between early DCIS and invasive cancer regions (PPLR after Bonferroni correction < 0.01) (**Fig. 2D-E, Table S6**). For most genes the expression level is higher in the invasive cancer (**Fig. S4B)**. The expression signal density in late DCIS is usually somewhere between early DCIS and invasive cancer, although differences exist between genes. Most immune panel genes behave in a similar fashion, exhibiting less than 25% of the signal enrichment detected in the invasive cancer (**Fig. 3D-E)**. This is consistent with the picture whereby immune cells, by and large, remain outside of the ducts irrespective of genetic features of the clone inside (**Fig. 3A**, box b; **Fig. S4C**). In late DCIS, some immune cell signals appear clustered close to ‘microbreaches’ in the mesenchymal cell layer (arrowheads, box c; **Fig. S4C**). Interestingly, in one region, immune signals cluster specifically with a group of purple clone cells in a duct that is otherwise occupied by an evolutionarily distant orange clone – a picture that, in the absence of the precursor DCIS clone, might support a pattern of cancer reinvasion (far right box; **Fig. 3C, Fig. S4D**). A smaller group of immune marker genes including T-regulatory cell marker (*TNFRSF18*) and the mainly cancer cell derived MHC class I chain related-proteins A and B (*MICA, MICB*) do not exhibit clustering but are diffusely scattered throughout the purple DCIS. This pattern is also apparent in the pure DCIS comparisons (orange versus green) in the previous section, it is therefore conceivable that these more diffuse immune changes might gradually emerge during waves of clonal progression (**Fig. 2E-F**).

The oncology panel genes are more heterogeneous in their expression behaviour and this undoubtedly reflects the more diverse composition of the panel. For around half of the genes, at least 50% of the increase in expression level seen in the invasive cancer is evident within the late DCIS (left box; **Fig. 3A, Fig. 3D-E**). A range of genes, including prognostic genes linked to both proliferation (*Ki67*) and invasion (*CTSL2*), have similar expression levels in late DCIS and invasive clones (**Fig. 3E**)(*32*). These gene signals are diffusely distributed throughout the purple clone regions without clustering indicating they are a general property of the DCIS cells. It therefore appears that at the DCIS stage, neoplastic cells can be armed with both the genetic and many of the transcriptional changes needed for life as an invasive cancer. It is perhaps remarkable that these cells form extensive expansions within intact ducts rather than invading immediately. While these data support the model of intraductal clonal progression they also indicate that the ultimate invasive transition could well be driven by the tumour microenvironment or rapid expansion of one clone could physically rupture the ductal basement membrane, spilling multiple clones into the breast stroma (*21, 30*).

### Tracing the emergence of aggressive cancer characteristics in a lymph node metastasis

Lymph node metastasis predicts distant metastasis and death, but whether it plays an active role in facilitating cancer progression or simply reflects more aggressive primary cancer biology is unknown. To assess whether the lymph node might drive clinically meaningful evolutionary progression, we selected a case where the clinically targetable breast cancer oncogene *HER2* was found to be amplified in the lymph node but not primary tumours by WGS (*9*) (**Fig. 4A, Fig. S5A**). As in case 1, we designed BaSISS padlock probes to mutations from the branches of the WGS inferred tree (stars; **Fig. 4A**). Targets included a passenger mutation in *HER2* and a novel internal ‘fusion’ in *CACNB1* (a gene 5 prime to *HER2*). These mutations are predicted to have occurred prior to and during the breakage fusion bridge event that generated the *HER2* amplification event respectively, and were included to aid evolutionary timing (**Fig. 4A, Fig. S5A, Table S1)**.

The BaSISS signal data exhibit spatial patterns that support the existence of at least 2 lymph node clones: A post-amplification clone with high *HER2* mutation density and *CACNB1* signals and at least one pre-amplification clone with low level *HER2* mutation signals but similar trunk mutation density (BaSISS plots; **Fig. 4B-C**). This supports WGS subclonal copy number data (**Fig. S5A**). We therefore provided the model with a genotype matrix for the 4 clones indicated in **Figure 4A** (**Table S1**), generating the clone maps of primary cancers and the lymph node in **Figures 4D-E**.

A subclonal population (purple) is detected in one of the primary cancers and this exhibits histological and expression features of more aggressive disease compared to the more ancestral (blue) clone (**Fig. S4B-D**). However, we did not find any evidence to support the emergence of the *HER2* amplified (orange) clone or the *HER2* mutation bearing, pre-amplification (green) clone within the primary cancers here or on a serial tissue section. Nonetheless, the existence of multiple clones in the lymph node confirms that *HER2* amplification was not necessary to establish metastasis in this cancer. Although we cannot exclude the possibility that *HER2* amplification occurred in an unsampled region of the primary tumour, the presence of the green *HER2* mutant, pre-amplification clone in the lymph node is consistent with emergence at this site (**Fig. 4B-C**).

The lymph node contains more than one pattern of cancer growth and these patterns are associated with different genetic subclones. The first, a solid growth pattern composed of pure cancer cells with no obvious stromal component and a paucity of immune cells is entirely formed by the orange clone (**Fig. 4C, Fig. 4E)**. Orange deposits are surrounded by endothelial cell layers and neat lines of *CD34* and *PDGFRB* ISS signals that are expressed by lymph node sinuses and potentially microvessels (**Fig. 4G, Fig. S5E)**. The remainder of the node is heavily infiltrated by cancer cells that exhibit a diffuse infiltrative pattern, intermingling with immune cells and frequently forming perifollicular aggregates (**Fig. 4C, Fig. S5F**). Most of these cells are assigned to one or other pre-*HER2*-amplification clone, although confidence of exact clone assignment is low due to low cancer purity in these areas. We therefore observe that the lymph node offers a distinct environment that permits segregated growth of genetically distinct subclones.

Consistent with a more aggressive emerging subclone, the orange subclone expresses higher levels of invasion and proliferation related genes, relative to the green and blue subclones (*32*) (**Fig. 4F; Supp. Methods**). Interestingly, three of the most highly enriched genes – Vascular endothelial growth factor A (*VEGFA)* (2.4 fold), carbonic anhydrase IX (*CA9*)(236 fold) and C-X-C motif chemokine ligand 8 (*CXCL8*)(384 fold) – are implicated in hypoxia and angiogenesis and their ISS signals follow striking patterns (PPLR =0.001) *(44*-*46*) (**Fig. 4G-H**). The signals are highly specific to the orange subclone and their acquisition is cumulative, following metastatic deposit size. For each gene, density within a deposit is inversely related to the distance from the endothelium and presumably the inverse oxygen gradient (**Fig. 4G)**. While *CA9* and *VEGFA* seem to be mainly cancer cell derived, *CXCL8* co-localises with CD45+ immune/lymphoid cells (by IHC) and *CD68/CD163* signals (by ISS) consistent with a macrophage origin in this scenario (**Fig. S4G)**. Both *HER2* amplification and hypoxic signatures are associated with adverse clinical outcomes in breast cancer (*47*). This case provides another powerful example of how spatial genomics can uncover the interplay of genetics and tumour microenvironment in contributing towards the subclonal diversification that underpins the emergence of clinically aggressive cancers.

While the eight samples interrogated with BaSISS revealed a wealth of information related to their individual patterns of subclonal growth, subclone-specific phenotypes and successive transcriptomic changes acquired during progression, some overarching patterns also emerge. Evaluating the clinical recurrence assay gene groups based on the prognostic ISS marker panel, we observe that proliferative and invasive group gene scores steadily increase along seven predicted clonal succession episodes spanning three cancer stages from DCIS to lymph node metastasis (**Fig. S6**). This would be consistent with the notion that major clonal succession events in a cancer reflect the emergence of increasingly proliferative clones (*48*). Larger spatial genomics studies may guide the development of precise biomarkers and molecular staging systems by exploiting the sequential nature of subclone specific changes.

## Discussion

Here we present BaSISS, a highly multiplexed fluorescence microscopy based protocol to map and phenotypically characterise cancer clones. The BaSISS technology supports a tailored approach according to each cancer’s unique complement of mutations. In this proof of principle study we map relatively broad subclone populations identified through multi-region WGS but the spatial genomics approach could equally be applied to more detailed phylogenies. A particular advantage of the technology is that it is capable of interrogating very large tissue sections and that it is comparably cheap, unlike solely relying on sequencing based methods (*49*). In theory, the approach holds the potential to create three dimensional genomic tomographs by aligning consecutive tissue sections. A limitation of the approach is relatively low sensitivity, which currently precludes single cell genotyping. For standard ISS, additional sensitivity can be achieved by tiling transcripts with more probes; unfortunately, this is not feasible for point mutations at a defined genomic location. A switch to hybridisation based sequencing and direct RNA binding probes, which eliminate the requirement for reverse transcription are currently limited to gene expression, but with further development should also improve base specific detection several fold (*50, 51*).

BaSISS’ ability to spatially locate and molecularly characterise different cancer subclones adds essential features to the spatial genomics tool kit. It provides a robust evolutionary framework that is necessary to interpret the biological relevance of many of the more plastic spatial characteristics of a cancer. Future widespread application of spatial genomics approaches will uncover how cancers grow in different tissues and allow us to track, trace and characterise the ill-fated clones that are responsible for adverse clinical outcomes.

## Supporting information

Table S1 probe design

Table S2 clinical

Table S3 bulk sequence data

Table S4 BaSISS target coverage

Table S5 ISS gene coverage

Table S6 clone gene expression

## Author contributions

JS, PJC, MN and LRY designed the study. AL, MG and LRY analysed and interpreted data and drafted the article and figures. AL and MG developed the core mathematical models. AS and VK contributed to mathematical modeling. AS contributed to image segmentation and processing. JS, MN and CS acquired ISS data and contributed to data interpretation and manuscript preparation. AR provided samples. ALR, LM, SP contributed histopathological expertise. YSJ contributed RNAseq expertise. SD contributed to WGS sub clonality analysis. JSP, VV, TL, OAB, MGataric contributed to development of bespoke ISS analysis pipelines. All authors reviewed and commented on the manuscript.

## Data and materials availability

Code that was used in the analysis could be found in https://github.com/gerstung-lab/BaSISS. Code used to segment nuclei in images is available in https://github.com/yozhikoff/segmentation. Data is available upon request.

## List of Supplementary Materials

Materials and Methods Fig. S1-S6

Table S1-S6

Methods references (*7, 9, 15, 28, 32, 33, 52*–*65*)

