## Supplementary Materials for "Spatial genomics maps the structure, character and evolution of cancer clones"

**This PDF file includes:**

Materials and Methods
Figs. S1 to S6
Captions for Tables S1 to S6

**Other Supplementary Materials for this manuscript include the following:**

Tables S1 to S6

### Materials and Methods

### 1 Tissue samples

Breast tissue samples were obtained from mastectomies performed for the diagnosis of multifocal primary breast cancer. Samples and data were obtained and managed in line with the declaration of Helsinki under protocol 93-085 (Dana Farber Cancer Institute). Sample and data handling at the Wellcome Sanger Institute, Cambridgeshire, UK was performed under the wider framework and approval for the Breast Cancer Genome Analyses for the International Cancer Genome Consortium Working Group under REC reference: 09/H0306/36.

Tissue blocks were first sliced to obtain sufficient nucleic acids for bulk whole genome sequencing (WGS) and RNAseq analysis. Subsequent serial 10um sections were obtained from the same tissue block and orientation for base specific in-situ sequencing (BaSISS), in-situ sequencing (ISS) and histological assessment.

### 2 Pathological assessment

For each tissue block a frozen section was prepared for pathological review using standard haematoxylin and eosin (H&E) staining. H&E review and annotatation was performed by experienced breast histopathologists.

For immunohistochemical staining after ISS, cover glasses were detached by over-night incubation in TBS-Tween20 (0.05%). Slides were fixed for 10 min with 3.7% formaldehyde (Sigma Aldrich, Munich, Germany). Endogenous peroxidases were blocked with DAKO REAL peroxidase-blocking solution (Agilent, Glostrup, Danmark) for 10 min at room temperature, followed by incubation with DAKO serum-free protein blocking solution (Agilent) for 30 min. Sections were incubated with mouse monoclonal PanCK antibody (clone AE1/AE3, Agilent) diluted 1:100 in DAKO REAL antibody diluent (Agilent) overnight at 4°C in an humidity chamber. Thereafter, secondary ImmPRESS HRP Anti-mouse IgG (VectorLaboratories, Orton Southgate Peterborough, UK) reagent was applied for 30min and chromogenic visualization was performed with DAB Peroxidase substrate kit (Vector laboratories) according to the manufacturer's instructions. Slides were counterstained with Mayer's HTX Hematoxylin (Histolab, Gothenborg, Sweden) for 30 seconds, dehydrated and permanently mounted (Vecta Mount, Vector laboratories).

### 3 Inferring subclone composition and evolutionary histories from bulk genomic data

The subclone composition estimates from multi-region WGS somatic substitution and copy number data used in this analysis are those reported in the original publications *(7, 9)*. The approach detects a finite number of subclones each with defined parameters including the number of mutations within that subclone and the fraction of cells from each sample that contain those subclone specific mutations – the results for each sample are summarised in **Table S2**.

#### 3.1 Bulk tissue sequencing

To generate DNA and RNA sequence data 10 x 10-20um serial slices of tissue blocks were pooled and homogenised followed by nucleic acid extraction. Short insert whole genome and targeted capture paired-end libraries and 350bp poly-A selected RNA libraries were created, flow cells prepared and sequencing clusters generated according to Illumina protocols *(53)*. 100 base whole genome sequence data, 150 base targeted capture genomic data and 75 base RNA sequence data was generated using Illumina HiSeq 2000® genome analyzers. Genomic and RNA sequence data were mapped to the reference human genome (GRCh37 Ensembl 58) using Burrows Wheeler Aligner

*(55)* and TopHat (v1.3.3), respectively. Original data associated with the prior publication datasets from which these samples derive are deposited in the European Genome Phenome Archive (EGA) with the following accessions: EGAD00001002696 and EGAD00001000898. Bulk sequence data metrics are reported in **Table S2**.

#### 3.2 Somatic mutation identification

Genome-wide somatic substitutions were called using CaVEMan (Cancer Variants Through Expectation Maximization: CaVEMan). Rearrangements were identified from discordantly mapping reads using BRASSII (BReakpoint AnalySiS). Data is available for download from `sftpsrv.sanger.ac.uk/SpatialGenomics`.

#### 3.3 Cancer cell fractions and mutation clustering

Subclonal clusters anad clustering approaches were as previously reported in the original publications *(7,9)* and full methodology is described in *(52, 56)*. Briefly, variants were clustered into groups of consistent cancer cell fractions, based on the number of supporting reads, underlying copy number and cancer purity. The most up-to-date version of the model, known as 'DPClust' is available here: Wedge-Oxford/dpclust.

#### 3.4 Principles of Phylogenetic Tree Construction

To infer the evolutionary relationship between subclones, phylogenetic trees were constructed using the "pigeon hole principle" which states that if there are $n$ objects that are placed in $m$ containers if $n > m$ then at least one container must contain more than one object. Extending this to subclonal reconstruction we can appreciate that the sum of subclone CCFs cannot exceed the CCF of their ancestor. For example, if there are $3$ subclones with CCFs of $1$, $0.9$ and $0.5$ then there must be a linear evolution – as $1+0.9 > 1$, CCF subclone $0.9$ must be a descendant of the CCF subclone

1; furthermore, as $0.9 + 0.5 > 1$, CCF subclone $0.5$ must be a descendant of the CCF subclone $0.9$.

In contrast one may see the possibility of branching evolution where the sum of subclone populations is less than $1$. For example, $3$ subclones with CCFs of $1$, $0.5$ and $0.4$ could arise either through linear evolution: $1 > 0.5 > 0.4$ or along separate ancestral lines because $0.5 + 0.4 < 1$. When applying these principles across multi-region samples the compatible underlying tree structures is usually greatly restricted because of the requirement that the same tree structure must be compatible across all samples. Applying these principles one can identify the phylogenetic tree structure(s) most compatible with the underlying data. Final matrices associated to the derived clonal phylogenetic trees are presented in **Table S3**.

### 4 BaSISS and ISS protocols

#### 4.1 Subclone specific somatic mutation target selection

In case PD9694, WGS identified 3,372 somatic substitutions across samples PD9694a,c and d. Applying the mutation clustering approach as described above, to these data identified a total of 8 clusters, 6 of which were considered robust and retained (they contained >1% of all mutations) **Table S2**. Individual clusters contained 163-817 mutations. Higher depth targeted capture resequencing (average of 150X) of 317 somatic mutations from each subclone was performed. 100% of somatic mutations were validated and applying the mutation clustering approach to this independent dataset reiterated subclone properties identified in the original experiment (**Table S2**). Applying the pigeonhole principle to the data identified 2 trees that were most compatible with the data, and that differed from each other according to the position of one branch (green) as depicted in **Fig S2E**.

To map subclone architecture within the tissue space, and to see if spatial subclone segregation patterns may help to resolve the tree, we identified somatic mutations from each subclone that would be used for padlock probe design. We selected a total of 25 coding mutations: 11 that reported the clonal cluster (cluster 8, grey) and 11 that reported subclonal clusters (6 from cluster 2, purple; 2 from cluster 3, blue; 1 from cluster 6, red, 1 from cluster 7, yellow and 1 from cluster 1, green) and 3 that reported on mutations that were present in a subset of samples but not assigned to a cluster cluster because they fell in regions of differential copy number. Two probes were also designed to detect *FGFR1* amplification (correlates with yellow cluster) (**Table S3**).

In case PD14780, WGS identified 7,772 somatic substitutions, 162 of which occurred in coding regions. In this experiment we used RNAseq to identify which mutations were expressed and provided confirmatory evidence for patterns of expression across the related samples. We identified 7 mutations that had more than 20X coverage using BWA or Tophat. We designed probes for 3 mutations that were proven to be expressed and showed consistent patterns of heterogeneity in RNAseq and WGS data – for the first mutation (*RPL37A*:r.985g>c) all but 2 reads reporting the mutation were detected in sample PD14780a, for the second mutation (*ERBB2*:r.4370g>u) all but 1 read reporting it was limited to PD14780e and for the final sample (*COX19*:r.394c>g) all samples expressed the mutation. We also designed a probe for a mutation (*PLXNA1*:r.7031c>u) that was expressed in all samples but for one sample (PD14780e) there was no supporting genomic evidence despite 30X coverage at the mutation locus. This discrepancy could be a play of chance as this sample (a lymph node) had a very low predicted tumour purity (10%) or could reflect spatial segregation of a subclone containing this mutation in the lymph node as the RNA and DNA libraries are prepared from independent tissue homogenates created from serial sections and so may not be perfect biological replicates. The ISS experiment was intended to differentiate between these scenarios. We designed a probe for *TP53*:p.E180* that was determined to be a clonal driver mutation but was not determined to be expressed based on RNAseq to determine if low/ absent expression of a mutation negates its detection in ISS data.

We also designed padlock probes towards a sense anti-sense fusion transcript identified in *CACNB1*. The average gene expression level of this gene was identified to be elevated in sample PD14780e compared to sample D14780a and D14780d (FPKM = 4.4 versus 2.13 and 1.17). Localisation of the gene identified that an inversion rearrangement involving introns 9 and 12 resulting in a novel splice site in antisense intron 10 resulting in antisense transcription of a novel exon formed by part of intron 10 and exon 10.

Exon 9 sequence:
ACAGAGCATGTGCCCCCCTATGACGTGGTGCCTTCCATGAGGCCCATCATCCTGGTG
GGACCGTCGCTCAAGGGCTACGAG

Antisense intron 10 part:
TCCGAGTCCCAGGATTGTGATTCGAGGCATCCTGCCCATCCCCAGCAGCAGAGAGCA
ACGGCAGGTGCGAGGAGCAGCTCCCAGGATCTTAC

Antisense exon 10 part:
CTGCCATCAAACCGATGCTTCAAGAAGTCAAATAAAGCTT

We designed padlock probes that spanned the antisense intron and exon parts of the novel exon 10 sequence (CACNB10-10 probe) and one that read through from exon 9 (3' arm) to the antisense intron 10 portion (5' arm). A probe was also designed to detect the normal exon splicing across exon 9-10 (CACNB9-10wt). Notably given the position of the gene within a series of breakage fusion bridges (that resulted in high level amplification of *ERBB2*) we expected both the mutant and wild type transcript to be more highly expressed in cells that had undergone this process.

Probes designed against 2 fusion genes predicted by an early implementation of deFuse (CARTPT:PID1,

ZNF652:SNHG5) were excluded from our downstream analysis as they were uninformative (Table S3). BaSISS signals did not faithfully localise to cancer cell populations. Further investigation revealed a lack of predicted fusion-gene/probe mapping specificity: with the ZNF652:SNHG5 probe mapping to the *SNHG5* wildtype sequence and the CARTPT:PID1 probe mapping to repetitive sequences and a histone mark.

#### 4.2 In-Situ Sequencing

##### 4.2.1 Tissue specimens

Serial 10um tissue sections were cut from fresh frozen tumour tissue blocks mounted on superfrost slides and sent from DFCI to ScilifeLab Stockholm, Sweden where ISS and IHC was performed under Karolinska Institutes rules for the handling of blood and other human sample material, reference number 1-31/2019 (with a local HUMRA risk assessment form). Tissue blocks included the same ones used for the WGS and RNAseq experiments and additional tumour blocks were identified where applicable.

##### 4.2.2 Padlock probe design

The padlock probe consists of a single stranded DNA oligo with two target recognition arms in the 3' and 5' end of the oligo, enabling circularization of the padlock probe upon target hybridization. Each target recognition arm typically has a length of 15-25 nucleotides. The two arms are interspaced with a linker sequence comprising a 20 nucleotide anchor primer sequence and a target-specific 4 nucleotide barcode followed by a 5 nucleotide stabilizing sequence for sequencing-by-ligation. The used barcodes were selected in such a way that they differ in at least two positions. In total, three padlock probe gene panels were used, marker gene-, mutation- (one specifically designed for each case) and immune- expression panels. Padlock probes were ordered as ultramer DNA oligos from Integrated DNA Technologies (IDT, Leuven, Belgium) with 5'-phosphorylation modification and were reconstituted in Tris/EDTA buffer.

###### 4.2.3 Mutation panel designs

Two separate case-specific mutation panels were designed, the selection of gene targets is mentioned above. For wildtype- and mutation-specific padlock probes, the sequence upstream of the mutated site resembles the 3' arm of the padlock probe, with the wildtype or mutant nucleotide at the 3' end, whereas the sequence downstream of the mutated site resembles the 5' arm. The target recognition arm lengths were adjusted to have a similar melting temperature of $\sim 55°C$. For each mutation site, one wild-type and one mutation-specific padlock probe was designed. For case PD9694, in addition to the anchor and barcode sequences a 20 nucleotide sequence used for a hybridization cycle was added in the padlock probe linker sequence. The padlock probe sequences of the two mutation panels are listed in **Table S3**. Specific primers were used for the in situ reverse transcription and were designed as the reverse complement sequence of the 5' target recognition arm. Primers were ordered as DNA oligos from Integrated DNA Technologies (IDT) and were reconstituted in Tris/EDTA buffer.

###### 4.2.4 Immune panel design

Large scale probe design was facilitated using an in-house Python software package as described previously (Ref PMID 31740815) which utilizes ClustalW and BLAST+ to ensure probe specificity. Each padlock probe of the immune panel was designed to contain two 20 nucleotide long target recognition arms. Only target fragments with melting temperature between 65°C and 75°C were considered. Probes were selected aiming to obtain a distribution along the whole length of the transcript.

Overall five padlock probes were selected per target gene with exception of *HLA-DRB1*, where only two specific probes could be designed. For the T cell specific genes *CD8A, FOXP3, EOMES, CD4* and *IFNG* 20 probes were selected in order to increase the detection efficiency for these target genes. All padlock probe sequences of the immune panel are shown in **Table S3**.

A combination of random decamer primers (IDT, Leuven Belgium) and specific primers were used for in situ reverse transcription. Specific primers were designed to hybridize to the mRNA 15-20 nucleotides downstream of the target sequence. Primers were ordered as DNA oligos from Integrated DNA Technologies (IDT) and were reconstituted in Tris/EDTA buffer.

Target genes for the immune panel were selected to cover a broad range of immune cell subtype marker with special emphasis on T cell subsets and their regulation:

| | | | | | |
|---|---|---|---|---|---|
| Pan leukocyte marker | *CD45* | | | | |
| General T cell marker | *CD3D* | *CD4* | *CD8A* | *CD8B* | *CD45RO* |
| | *CD45RO* | *TNFRSF4* | *TBX21* | *EOMES* | *BTLA4* |
| Activated T cells | *CD25* | *PRF1* | *IFNG* | *GZMB* | *IL2* |
| | *MKI67* | *FASLG* | *TNFRSF9* | *ICOS* | |
| Regulatory T cells | *FOXP3* | | | | |
| Checkpoint Inhibition | *CD274* | *PDCD1* | *LAG3* | *CTLA4* | *HAVCR2* |
| | *VSIR* | *TNFRSF18* | *KLRG1* | | |
| Regulatory factors | *CCR7* | *IL7R* | *IL4* | *IL5* | *IL10* |
| Immunevasion | *TGFB1* | *FAP* | | | |
| NK cells | *NCR1* | *MICA* | *MICB* | *NCAM1* | *NKG2D* |
| Macrophages | *CD68* | *CD163* | *CD83* | *CD80* | *CD86* |
| | *CD14* | | | | |
| Macrophages | *CXCL8* | *ENTPD1* | *FCGR3* | *FUT4* | |
| Dendritic cells | *ITGAM* | *ITGAX* | | | |
| B cells | *MS4A1* | | | | |
| Tumor environment | *CSF1R* | *CD34* | *ARG1* | *ARG2* | *IDO1* |
| | *IDO2* | *TDO2* | *NOS2* | *HLA-DR1* | *CA9* |

###### 4.2.5 Oncology gene panel design

The oncology gene panel includes genes included involved in proliferation, EMT, invasiveness, stemness, angiogenesis as well as genes for breast cancer subtyping and oncotypeDX recurrence scoring *(*28, 32). The target recognition arms were designed to capture most splice variants of the gene transcripts and blasted to confirm their specificity. Each target recognition sequence had a GC content of 50-55% and a melting temperature of $\sim 55°$C. In this older design, the panel included one padlock probe per gene target and the barcodes used were only differing in one position. For the marker gene panel, random decamer primers were used for *in-situ* reverse transcription (IDT, Leuven Belgium).

###### 4.2.6 *In-situ* sequencing (ISS)

ISS was performed as described *(15)*, and modified according to *(28)*, and was used to spatially resolve oncology panel-, mutation panel- and immune panel- gene expression profiles on consecutive sections from the breast tumor tissue blocks. The ISS library preparation and sequencing is described in detail at protocols.io.bb2giqbw, the steps in the protocol that were modified for breast cancer tissues are indicated below. In brief, library preparation included fixation of tissue sections with 4% PFA for 30 min (step 2) followed by permeabilized with 0.1 mg/ml pepsin (Sigma) in 0.1 M HCl 37°C for 90 s (step 4). SecureSeal$^{TM}$ reaction chambers were mounted on top of the tissues (Grace Biolabs, Bend, United States) and cDNA was synthesized insitu using specific DNA primers (125nM for mutation panel, 5nM for immune panel) and/or random decamer primers (5$\mu$M for immune and marker gene panels) (step 10) (IDT, Leuven Belgium, sequences are listed in **Table S3**). Rnase H was used to generate single-stranded cDNA that the padlock probes could hybridize to. Hybridized padlock probes (10nM of each in immune panel, 0nM of each in mutation and marker gene panel, step 15) (IDT, Leuven Belgium, sequences are listed in **Table S3**) were ligated using Tth ligase, a highly specific DNA ligase that can discriminate correct base-pairing at the single nucleotide level. Only completely target-complementary padlock probes become ligated, forming closed circles that could then be amplified through rolling circle amplification (RCA). Of note, for case PD9694, the experimental conditions for the first replica of ISS differed slightly with a diverse Phi29 buffer (Thermo Fisher 10X reaction buffer: 330 mM Tris-acetate (pH 7.9 at 37 °C), 100 mM Mg-acetate, 660 mM K-acetate, 1% Tween 20 and 10 mM DTT) and no Exonuclease 1 in the rolling circle amplification step

The four-five nucleotide target-specific barcodes included in the padlock probe linker sequence were clonally amplified in the RCA products allowing identification through sequencing by ligation (according to Dermat et al) of anchor primer and fluorophore-labelled interrogation probes *(28)*. Nuclei were stained with 4',6-diamidino-2-phenylindole (DAPI). The target-specific barcodes were sequenced with four sequencing and imaging rounds. For case PD9694, the mutation panel was sequenced with one hybridization cycle in addition to the four sequencing by ligation cycles. After the ISS analysis, the tissue sections were stained with PanCK, CD24 or KI67 antibody.

###### 4.2.7 Imaging

Images were acquired with an automated Zeiss Axioplan II epifluorescence microscope (Zeiss, Oberkochen, Germany) using a z-stack of $0.49\mu\text{m}\times 11$ and a tile overlap of 10%. Images were scanned with a 20× objective. For the first base sequenced, the exposure times were calibrated so that the signal intensity values were similar for all sequencing channels (A–Cy5, G-Cy3, C-Texas Red and T-AF488), the calibrated exposure times were then kept constant for all remaining sequencing cycles. Orthogonal projections and stitching of tiles were done with the ZEN software (Zeiss).

### 5 ISS signal localisation and deconvolution

#### 5.1 Image stitching

From the total of 51 image sets, 43 were stitched with Carl-Zeiss ZEN software (version 3.1), and the other 8 failed image sets were stitched using BigStitcher (version 0.9) *(62)*.

#### 5.2 Image registration

The registration across imaging cycles was performed in two steps: affine registration on DAPI channel and subsequently local warping on anchor channel. For both steps we used algorithms provided in libraries OpenCV-contrib (version 4.3.0) *(60)* and scikit-image (version 0.17) *(61)*. In all imaging cycles, before the registration, both DAPI and anchor channels were maximum intensity projected across the image z-stack.

During the affine registration step, we coarsely align images of all cycles to the first one based on the DAPI channel. Firstly, we detect key points in the images of each cycle using the FAST feature detector. Secondly, for each key point, its surrounding area is described with histograms of oriented gradients using the DAISY feature descriptor. After that, using the key points and their descriptors, the FLANN-based matcher finds correspondences between pairs of key points from reference and moving images and filters out unreliable points. Lastly, the remaining key points are processed using the RANSAC-based algorithm that aligns them and estimates affine transformation parameters with 4 degrees of freedom.

The second registration step aligns imaging cycles sequentially using the anchor channel (fluorophore Cy7). We applied Farneback optical flow algorithm to achieve more accurate registration by warping the images locally, so that RNA spots of different channels can be better aligned despite the presence of nuclei swelling, imperfect stitching and sample distortion.

In both steps, we optimized the algorithm by performing computation on the tiled images to reduce memory consumption and accelerate the transformation parameters estimation.

#### 5.3 Serial tissue image alignment

The dataset we constructed contains a series of consecutive cuts per sample – at least one for BaSISS and 2 for oncology and immune panels of ISS. To be able to compare them these slides have to be aligned. We performed a spline-based elastic registration implemented in ImageJ package UnwarpJ *(65)*. All ISS images were registered on the respective slide with BaSISS data.

#### 5.4 ISS signal deconvolution

After registration of images from different sequencing rounds, we locate RNA spots by applying the circular Hough transform to the reference anchor channel of the first round, which is implemented in MATLAB's function 'imfindcircles'. At the detected coordinates, image values are extracted from top-hat filtered coding channels across all sequencing rounds. We then perform decoding of the extracted image values via a Gaussian Mixture Model, where each mixture corresponds to one of the possible barcodes encoded via an experimental design. Finally, once the mixture model is fitted to the extracted image values, each detected spot is assigned to the most likely barcode. In addition to on-target barcodes, there is an infeasible class which represents RNA spots to which barcodes could not be assigned. In **Table S4** and **Table S5** we present QC metric of our datasets as a proportions of RNA spots with assigned barcodes ($p > 0.6$) to the total number of spots.

### 6 Nuclei segmentation

#### 6.1 Segmentation

A two-stage pipeline from Caicedo, Goodman et al. *(57)* was used for nuclei segmentation of a DAPI image. First, an ensemble of 32 pretrained neural networks was used to classify image pixels into three classes: background, nuclei and nuclear boundaries. The ensemble included neural networks with UNet- and FPN-like architectures with the following encoders: DPN-92, Resnet-152, InceptionResnetV2, Resnet-101. Then, the individual nuclei masks were obtained using the watershed algorithm. For each nucleus candidate, a set of morphological features (e.g. circularity, convexity, area, neighbours median area) was computed. After that, a gradient-boosting model (LGBM) based on these features was used to filter real nuclei from false-positive predictions. Code is available at yozhikoff/segmentation.

#### 6.2 IHC based classification

To quantitatively describe DAB-stained areas, Haematoxylin-Eosin-DAB colour convolution was used *(59)*. The implementation from the scikit-learn Python package was used.

### 7 Spatial mapping of cancer clones

Raw BaSISS data informs us of allelic variants spatial distribution. However, this data can not be directly interpreted in terms of cancer clones. To make sense of BaSISS data we designed a Bayesian model that aims to decompose local BaSISS signal counts into clone densities and corresponding genotypes.

#### 7.1 Core model

The BaSISS protocol records a series of fluorescent spots, which are decoded into a class of different barcodes $A$ corresponding to each targeted allele. This information produces a table of tuples $(a_i, x_i, y_i)$, where $a_i$ is the allele of spot $i$ and $x_i$ and $y_i$ are its respective two dimensional coordinates.

The counts of all probes can be represented by a three dimensional array $\mathbf{D} \in \mathbb{N}^{|a|\times|x|\times|y|}$, where $a$ refers to the allele and $x$ and $y$ are course grained coordinates on a regular grid of dimensions $|x| \times |y|$. The grid size was chosen to be 108.8$\mu$m considering a trade-off between data sparsity, precision and computational cost.

The essential idea is that the expected number of BaSISS signals is decomposed into maps of $|s|$ distinct clones $s$ $\mathbf{M} \in \mathbb{R}^{|s|\times|x|\times|y|}$ each with a distinct genotype $\mathbf{G} \in \mathbb{N}^{|a|\times|s|}$,

$$\mathbb{E}[\mathbf{D}] \approx \mathbf{G} \times \mathbf{M} = \sum_{s\in\text{subclones}} \mathbf{G}_{\cdot,s}\mathbf{M}_{s,\cdot,\cdot} \quad . \tag{1}$$

The genotype matrix $\mathbf{G}$ contains the number of each allelic copy in each clone $s$. $\mathbf{G}$ is a matrix representation of the underlying phylogenetic tree and provides the instructions of the allelic configuration in each branch of the tree.

The maps $\mathbf{M}$ provide the relative prevalence of each clone in a given area of the grid. As $\mathbf{M}$ should replace a spatially continuous map, it is modelled by softmax transformed two dimensional latent Gaussian processes.

#### 7.2 Latent Gaussian Process modeling of spatial clone maps

Depending on the choice of targeted alleles, signal counts in matrix $\mathbf{D}$ may appear rather sparse. For a robust clone prevalence map inference we use Gaussian processes as it allows to naturally utilise spatial distance information. Each clone $s$ is modeled as independent latent two dimensional Gaussian process with $\mu = 0$ and covariance matrix $K$ with RBF kernel over grid coordinates. Bandwidth $l$ was empirically chosen for sample PD9649 and adjusted for sample PD14780 according to the median signal/nuclei ratio.

$$K_x(x, x') = \exp\left[-\frac{(x - x')^2}{2l^2}\right] \tag{2}$$

$$m_s(x, y) \sim \mathcal{GP}(0, K_x \otimes K_y) \tag{3}$$

Each Gaussian process was mapped to simplex space using a softmax (multidimensional logistic) transformation with temperature parameter $\tau = 0.5$ which represents our weak prior belief that clones are intermixed.

$$\mathbf{M} = \left[\frac{e^{\tau m_0}}{1 + \sum_{i=0}^{|s|-1} e^{\tau m_i}}, \ldots, \frac{e^{\tau m_{|s|-1}}}{1 + \sum_{i=0}^{|s|-1} e^{\tau m_i}}\right] \tag{4}$$

##### 7.3 Sources of noise

To fit the above model to data a number of sources of noise need to be considered.

1. Cellular density $\boldsymbol{\nu}$
2. Differential probe specificity $\boldsymbol{\iota}$
3. Allelic confusion $\boldsymbol{\tau}$
4. Clone expression variations $\boldsymbol{\gamma}$
5. Homogeneous and inhomogeneous background adjustments $\boldsymbol{\beta}$
6. Overdispersed sampling fluctuations $\boldsymbol{\alpha}$

Accounting for these sources of noise, the equation for the expected number of BaSISS signals (1) $\boldsymbol{\mu} = \mathbb{E}[\mathbf{D}]$ becomes:

$$\boldsymbol{\mu}_{a,x,y} = \underbrace{\boldsymbol{\nu}_{x,y}}_{\text{cell density}} \cdot \overbrace{\boldsymbol{\iota}_a}^{\text{detection rate}} \cdot \underbrace{\sum_{a'} \boldsymbol{\tau}_{a,a'}}_{\text{probe confusion}} \sum_s \overbrace{\boldsymbol{\gamma}_{s,a}}^{\text{clone-specific expression}} \underbrace{\mathbf{G}_{a,s}\mathbf{M}_{s,x,y}}_{\text{clone contribution}} + \overbrace{\boldsymbol{\beta}_a}^{\text{background}} . \tag{5}$$

The exact nature of and statistical models for the different noise terms will be discussed in the next subsections.

###### 7.3.1 Cellular density

Rather than reporting abstract clone prevalence it's more insightful to bound them to actual cell densities. As a part of BaSISS signal deconvolution, DAPI stained image of nuclei is obtained. We use NN model to segment these images and store nuclei counts in a two dimensional array $\mathbf{N} \in \mathbb{N}^{|x| \times |y|}$. However, these counts might not always correctly represent cell densities due to possible segmentation errors and the fact that only a fraction of cell might be captured on the slide.

Therefore we treat nuclei counts $n_{x,y}$ as Poisson distributed values with mean $\boldsymbol{\nu}_{x,y}$ representing cell densities which in turn has a week Gamma prior.

$$\mathbf{N}_{x,y} \sim \text{Poisson}(\boldsymbol{\nu}_{x,y}) \tag{6}$$

$$\boldsymbol{\nu}_{x,y} \sim \text{Gamma}(\mu = 50, \sigma = 100) \tag{7}$$

###### 7.3.2 Differential probe specificity

BaSISS ability to detect allelic variants relies on a multistage process of reverse transcription, padlock probes annealing, amplification and fluorophore detection. As it is hard to take into account all possible sources of variability, we model variant detection rate $\boldsymbol{\iota} \in \mathbb{R}_{+}^{|a|}$ for each allele $a$.

$$\boldsymbol{\iota}_a \sim \text{Gamma}(\mu = 0.5, \sigma = 1) \tag{8}$$

###### 7.3.3 Allelic confusion

The difference between wild type and mutated allelic variants is often just a single nucleotide. This increases chances for the padlock probe to anneal to the wrong allelic variant resulting in allelic confusion. To model this confusion we design a sparse transition matrix $\boldsymbol{\tau} \in \mathbb{R}_{(0,1)}^{|a|,|a|}$ populated with transition probabilities $\{\boldsymbol{\tau}_{a,a'}\}$ for each allele. We put a strong regularising Beta prior on $t_{a,a}$ to shift probability density close to 1 and prevent it going below 0.75 to avoid non-identifiabilities.

$$\boldsymbol{\tau} = \left(\begin{array}{c|cccc} & \dots & wt & mut & \dots \\ \hline \vdots & & & & \\ wt & & \tau_{11} & 1-\tau_{11} & \\ mut & & 1-\tau_{22} & \tau_{22} & \\ \vdots & & & & \end{array}\right), \tag{9}$$

$$\boldsymbol{\tau}_{a,a'} \sim 1 - \text{Beta}(\alpha = 1, \beta = 25)/4$$

###### 7.3.4 Clone allelic variations

Although the mean clonal expression level is registered in detection rate matrix $\boldsymbol{\iota}$, it is reasonable to assume that different clones may have some level of diversity in expression of some genes. We model this deviation as a matrix $\boldsymbol{\gamma} \in \mathbb{R}_+^{|s|\times|a|}$ with a log-normally distributed prior such as $\mathbb{E}[\gamma] = 1$.

$$\boldsymbol{\gamma}_{s,a} \sim \mathrm{LogN}(\mu = 0, \sigma = 0.05) \tag{10}$$

###### 7.3.5 Homogeneous and inhomogeneous background adjustments

Raw BaSISS data comes from biological images which have additional sources of variability. The first simple one is the homogeneous additive shift $\boldsymbol{\beta} \in \mathbb{R}_+^{|a|}$ which increases expected value of a particular probe detection globally on the slide.

$$\boldsymbol{\beta}_a \sim \mathrm{Gamma}(\mu = 0.5, \sigma = 1) \tag{11}$$

The second one is inhomogeneous background shift which adjusts for local base signal detection variability. In practice it's hard to reliably model this type of background without a harsh regularisation due to it's flexibility. We encode inhomogeneous background shift as additional pseudo-clones $p$ which have corresponding relative prevalence $\mathbf{M}$ and pseudo-genotype $\boldsymbol{\Gamma} \in \mathbb{R}_+^{|p|\times|a|}$. Prevalence is modeled together with the real clones as a two dimensional latent Gaussian processes. Pseudo-genotype is sampled from a Beta prior which support is stretched to match median copy number $k$ among the loci. We use $|\psi| = 1$, but it can be increased for a more flexible background shift.

$$\boldsymbol{\Gamma}_{p,a} \sim \mathrm{Beta}(\alpha = 0.01, \beta = 1) \times k \tag{12}$$

###### 7.3.6 Sampling fluctuations

We model BaSISS counts as being Negative Binomial distributed given an unobserved mean detection level $\mu_{x,y,a}$ and over-dispersion parameter $\boldsymbol{\alpha}_a$ which accounts for unexplained variance:

$$\mathbf{D}_{x,y,a} \sim \mathrm{NB}(\mu_{x,y,a}, \alpha_a) \tag{13}$$

Over-dispersion $\alpha_a$ is sampled from Gamma distribution where distribution density is shifted towards larger values

$$\boldsymbol{\alpha}_a \sim \mathrm{Gamma}(\mu = 100, \sigma = 10) \tag{14}$$

##### 7.4 Optional inputs

Depending on which additional data accompanies investigated samples we may want to include them into proposed Bayesian framework. Currently, two option were used:

1. VAFs from adjacent slices
2. IHC based cell type counts

###### 7.4.1 VAFs from adjacent slices

WGS data contains information of the mutated allele frequencies on the whole slide. We can use this data to increase accuracy of our inference. To pass it into the model we calculate frequencies of each mutated variant $m$ on the slide $\mathbf{VAF}^{\text{BaSISS}} \in \mathbb{R}^{|m|}_{[0,1]}$ by summation of $\mathbf{M} \times \mathbf{G}$ product over spatial dimensions.

$$\mathbf{VAF}^{\text{BaSISS}} = \sum_{x,y} \mathbf{M} \times \frac{\mathbf{G}^{\text{mut}}}{\mathbf{G}^{\text{wt}} + \mathbf{G}^{\text{mut}}} \tag{15}$$

We then construct a Beta pseudo-likelihood of $\mathbf{VAF}^{\text{BaSISS}}$ with $\alpha$ and $\beta$ parameters proprtional to the number of mutated and wild-type reads in WGS experiment. We use parameter $u$ to inflate uncertainty in WGS data as it comes from proximal but not exactly the same slide we are observing in BaSISS experiment.

$$\text{VAF}_m^{\text{BaSISS}} \sim \text{Beta}(\alpha = \text{WGS}_m^{\text{mut}}/u + 1, \alpha = \text{WGS}_m^{\text{wt}}/u + 1) \tag{16}$$

To adjust for a disproportionately higher impact of spatial information on the total likelihood we multiply VAF log-likelihood by the number of tiles on the slide.

##### 7.4.2 IHC based cell type counts

IHC data reveals part of cell type heterogeneity. In our study we mainly worked with CD45 IHC staining which indicates location of lymphocytes. As we know that none of the lymphocytes should contribute to cancer clone weights, we could further regularise clone weights. This is done by construction of cell fraction matrix $\mathbf{CellFrac} \in \mathbb{R}_{[0,1]}^{|x| \times |y|}$ obtained by summation of clone map matrix $\mathbf{M}$ over clones of respected type $s^-$ or $s^+$.

$$\mathbf{CellFrac} = \frac{\sum_{s \in s^-} \mathbf{M}}{\sum_{s \in s^+} \mathbf{M}} \tag{17}$$

We use a Beta pseudo-likelihood of $\mathbf{CellFrac}$ distribution with $\alpha$ and $\beta$ parameters equal to cell counts of clear and IHC stained nuclei on the slide.

$$\mathbf{CellFrac}_{x,y} \sim \text{Beta}(\alpha = \text{IHC}^{-}_{x,y} + 1, \alpha = \text{IHC}^{+}_{x,y} + 1) \tag{18}$$

#### 7.5 Multi-sample version

In many cases some of the clones belonging to the same phylogenetic tree might not exist or exist only in a small proportion on most of the slides. Having a multi-sample generalisation of the proposed model helps us to correctly infer parameters associated to such clones. Essentially, the only parameters which are shared between all slides $k$ are: genotype matrix $\mathbf{G}$, clone allelic variations $\boldsymbol{\gamma}$, probe confusion transition matrix $\boldsymbol{\tau}$ and mean probe detection rate $\boldsymbol{\iota}$. All the other parameters become slide specific.

To allow some level of slide specific probe efficiency variation we multiply mean probe detection rate by a slide specific probe deviation matrix $\boldsymbol{\eta} \in \mathbb{R}_{+}^{|k|\times|a|}$ with a log-normally distributed prior such as $\mathbb{E}[\boldsymbol{\eta}] = 1$.

$$\boldsymbol{\eta}_{k,a} \sim \mathrm{LogN}(\mu = 0, \sigma = 0.05) \tag{19}$$

#### 7.6 Inference

Variational Bayesian Inference is used to approximate the posterior. We use mean-field version of Automatic Differentiation Variational Inference (ADVI) implementation from pymc3 package *(54)*. Briefly, within this framework, the posterior distribution over unknown parameters are approximated by appropriately transformed multivariate normal distributions with a diagonal covariance matrix. Inference is achieved by maximising log-likelihood of the data and minimising KL divergence from the posterior to prior, which are combined in the evidence lower bound (ELBO loss function).

Training is stopped when ELBO stops increasing passing at least 15,000 iterations of training using ADAM optimiser with learning rate 0.01. Posterior mean, standard deviation and quantiles for each parameter were computed using 300 samples from the variational posterior distribution.

#### 7.7 Evaluation of possible phylogenies based on BaSISS data

The phylogenetic tree solution constructed from multi-regional WGS data is often not unique. To further assist with the selection of the most likely tree among many, we could use BaSISS data to compute posterior odds ratios.

For example, if there are 2 possible phylogenetic trees with corresponding genotype matrices $\mathbf{G}_1$ and $\mathbf{G}_2$, we want to compute the posterior ratio of the genotype matrices given data. This could be expanded as a product of prior odds and of a ratio of marginal likelihoods.

$$\frac{P(\mathbf{G}_1|\mathbf{D})}{P(\mathbf{G}_2|\mathbf{D})} = \frac{P(\mathbf{G}_1)}{P(\mathbf{G}_2)} \frac{P(\mathbf{D}|\mathbf{G}_1)}{P(\mathbf{D}|\mathbf{G}_2)} \tag{20}$$

We estimate marginal log-likelihoods $\log P(\mathbf{D}|\mathbf{G})$ as ELBO obtained during model training.

### 8 Clone associated phenotype

Knowing spatial cell density distribution of clones we attempt to characterise the clones phenotypically. There are two characteristics we use – nuclei morphology and *in situ* sequencing expression signals. In both cases we characterise the whole region of high clone abundance. It is important to note that we do not distinguish cancer clone cells from any other type of cells and characterise selected location as a whole. The minimal unit of region is the grid tile.

For pure DCIS samples PD9694d and PD9694l and pure invasive sample PD14780a, we selected regions with the highest abundance of the respective clone. For samples with a mixture of DCIS and invasive morphologies – PD9694a, PD9694c – we have selected regions with the highest abundance of the respective clone within a border of a respective morphology (invasive or DCIS) as this was pertinent to the biological question. For a lymph node sample PD14780e, due to a large difference between solid and diffuse clones we considered regions in the top 20th percentile of the respective clone abundance on the whole slide. To avoid double assignment of regions we excluded intersecting ones.

#### 8.1 Expression association

##### 8.1.1 Core model

The essential idea behind the expression model is similar to the one we used in spatial mapping. ISS records a series of fluorescent spot, which are decoded into a set of barcodes corresponding to each of the targeted gene. Here we also represent counts of probes by a three dimensional array $\mathbf{D} \in \mathbb{N}^{|g|\times|x|\times|y|}$, where $g$ refers to the gene and $x$ and $y$ are coordinates on the grid.

The expected number of ISS signals is decomposed into maps of $s$ clone specific regions $\mathbf{M} \in \mathbb{R}_{\{0,1\}}^{|x|\times|y|\times|s|}$ each with a distinct expression pattern $\mathbf{F} \in \mathbb{R}_{+}^{|s|\times|g|}$,

$$\mathbb{E}[\mathbf{D}] \approx \mathbf{M} \times \mathbf{F} = \sum_{s \in \text{subclones}} \mathbf{G}_{\cdot,s}\mathbf{M}_{s,\cdot,\cdot} \quad . \tag{21}$$

However, here clone specific region matrix $\mathbf{M}$ is fixed. It is encoded as indicator function over grid tiles $i$ belonging to a region $R$ of respective clone $s$:

$$\mathbf{M}_{x,y,s} = \mathbf{1}_R(i_{x,y}) := \begin{cases} 1, & i_{x,y} \in R_s \\ 0, & i_{x,y} \notin R_s \end{cases} \tag{22}$$

Expression pattern $\mathbf{F}$ represents number of ISS signal for a particular gene $g$ per average cell in region belonging to clone $s$. We model it as a Gamma distribution

$$\mathbf{F}_{s,g} \sim \text{Gamma}(\mu = 0.5, \sigma = 1) \tag{23}$$

##### 8.1.2 Sources of noise

To fit the above model to data a number of sources of noise need to be considered.

1. Cellular density $\boldsymbol{\nu}$

2. Homogeneous background adjustments $\boldsymbol{\beta}$

3. Sampling fluctuations $\boldsymbol{\alpha}$

Accounting for these sources of noise, the equation for the expected number of BaSISS signals $\boldsymbol{\mu} = \mathbb{E}[\mathbf{D}]$ (21) becomes:

$$\boldsymbol{\mu}_{x,y,g} = \underbrace{\boldsymbol{\nu}_{x,y}}_{\text{cell density}} \cdot \underbrace{\sum_s \mathbf{M}_{x,y,s}\,\mathbf{F}_{s,g}}_{\text{clone regional expression}} + \underbrace{\boldsymbol{\beta}_a}_{\text{additive shift}} \tag{24}$$

The exact nature and statistical models for the listed noise terms is defined in exactly the same way as for BaSISS model, see sections 7.3.1, 7.3.5 and 7.3.6.

#### 8.2 Inference

Inference procedure is exactly the same as mentioned in section 7.6. Posterior mean, standard deviation, quantiles and other mentioned properties for each parameter were computed using 100.000 samples from the variational posterior distribution.

#### 8.3 Differential expression

Once a posterior distribution of clone associated expression $f_{s,g}$ is inferred, it is possible to compute the level an significance of differential expression between subclones. We use the probability of positive log-ratio (PPLR) which is defined as

$$P(\log f_{s_1,g} > \log f_{s_2,g}) = \int_0^{+\infty} P(\log f_{s_1,g} - \log f_{s_2,g})\,\mathrm{d}(\log f_{s_1,g} - \log f_{s_2,g}) \tag{25}$$

By setting a level of significance $\alpha$, this equation allows to find up-regulated genes associated with clone $s_1$ in comparison to $s_2$. To find down-regulated genes we just switch the clones and compare $s_2$ to $s_1$.

To control family-wise error rate we applied Bonferroni correction for each ISS panel and each direction of effect we testing to keep $\alpha = 0.01$.

#### 8.4 Clone associated nuclei phenotype

The following nuclei features were used to quantitatively describe the differences between cells of different subclones: intensity, diameter, roundness. Nucleus intensity was computed as the mean intensity of all pixels selected by applying the segmentation mask to the original DAPI image. To compute nucleus diameter and elongation, an ellipse was fitted to each segmentation mask. The roundness was defined as the ratio between the minor and the major axis of the fitted ellipse. Diameter was defined as the major axis of the ellipse.

#### 8.5 Clinical recurrence score groups

Due to the difference in expression ranges of ISS data and variability of reference genes such as *ACTB*, we thought that it is meaningless to compute recurrence score by the Recurrence-Score Algorithm *(32)*. However, it might be insightful to look at the relative group scores (GRB7, ER, invasion, proliferation) as they should indicate a direction of change.

Log transformed clone associated expression posteriors were used to compute group scores in accordance with the group formula from *(32)* and shifted to the positive domain by subtracting minimal value. PPLR was used to select the direction of significant transitions **Fig. S6**.

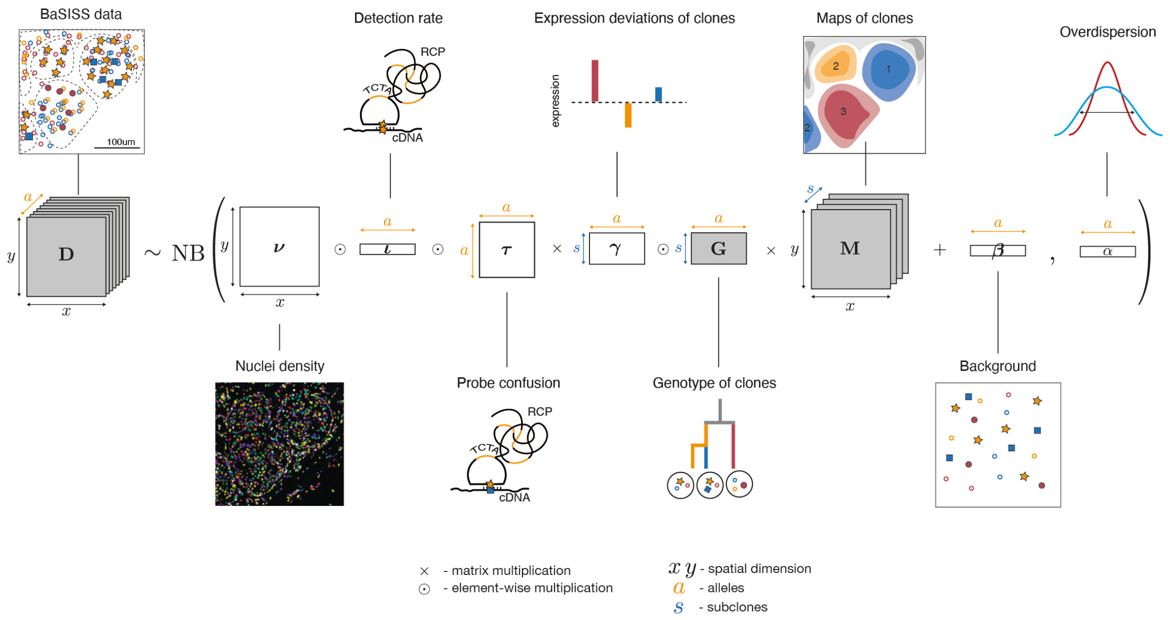

**Fig. S1 Mathematical modelling of spatial genomics data** Mathematical model for generating quantitative clone maps. The essential idea is that BaSISS signals count matrix D is decomposed into maps of clones M each with a distinct genotype G (grey shading), accounting for multiple sources of variability. For further details see Supplementary Methods section 7.

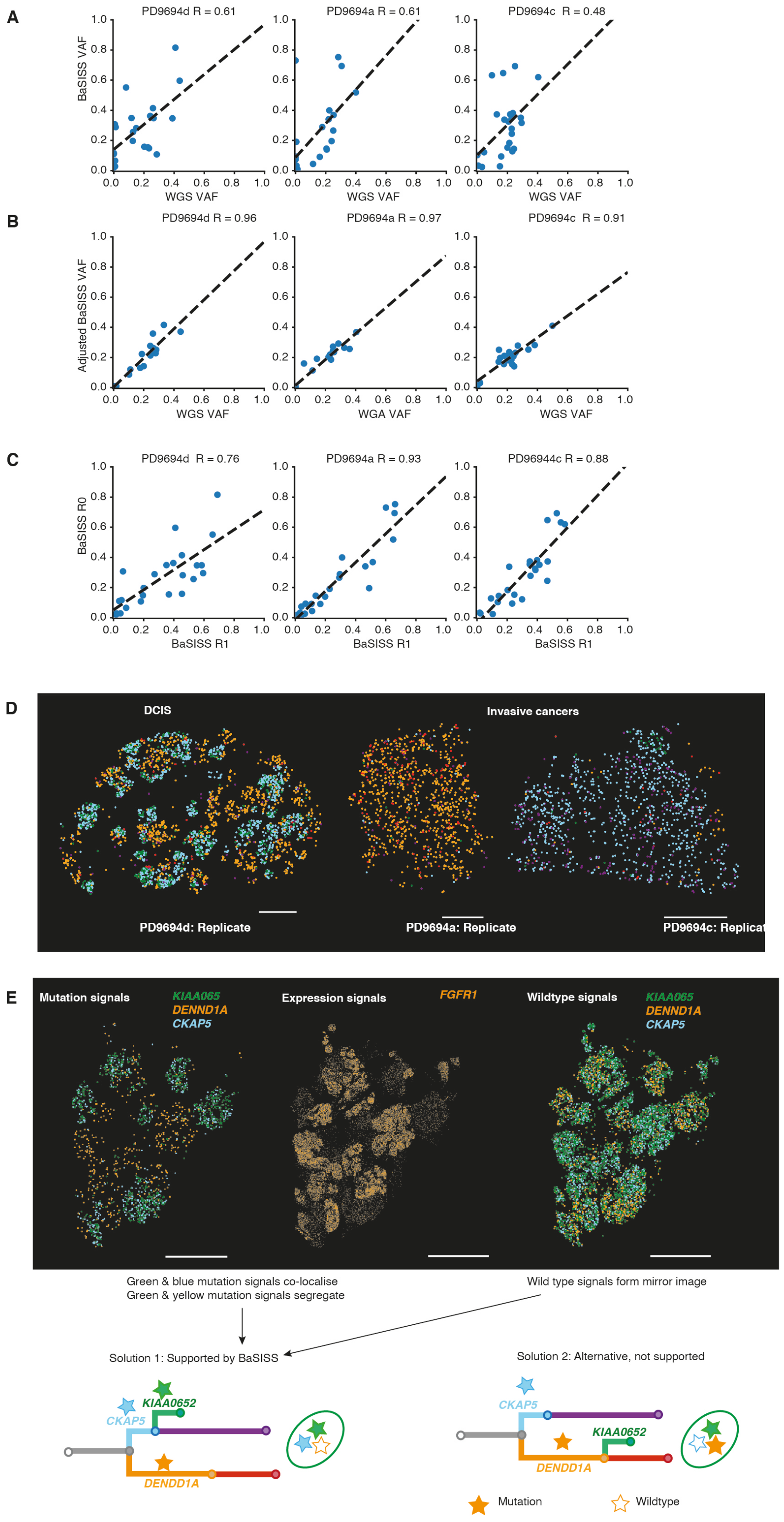

A
PD9694d R = 0.61
PD9694a R = 0.61
PD9694c R = 0.48
BaSISS VAF
WGS VAF
B
PD9694d R = 0.96
PD9694a R = 0.97
PD9694c R = 0.91
Adjusted BaSISS VAF
WGS VAF
WGA VAF
C
PD9694d R = 0.76
PD9694a R = 0.93
PD96944c R = 0.88
BaSISS R0
BaSISS R1
D
DCIS
Invasive cancers
PD9694d: Replicate
PD9694a: Replicate
PD9694c: Replicat
E
Mutation signals
KIAA065
DENND1A
CKAP5
Expression signals
FGFR1
Wildtype signals
KIAA065
DENND1A
CKAP5
Green & blue mutation signals co-localise
Green & yellow mutation signals segregate
Wild type signals form mirror image
Solution 1: Supported by BaSISS
Solution 2: Alternative, not supported
CKAP5
KIAA0652
DENND1A
CKAP5
KIAA0652
DENND1A
Mutation
Wildtype

**Fig. S2 Mapping cancer mutations in tissue context case PD9694** (**A-C**) Scatterplots of BaSISS variant allele frequencies (VAF) defined as number of mutation specific signals divided by mutation plus wildtype signals (depth) for each mutation target: (A) Compared to WGS VAF, (C) compared to WGS VAF after applying a probe specific adjustment (VAF)as described in methods section 7.4.1, (C) between replicate BaSISS experiments. R = Pearson's correlation coefficient. (**D**) BaSISS mutation signals for representative branch specific mutations (coloured according to the gene names that annotate branches of the tree in E) in a replicate experiment, corresponds to Figure 1C. (**E**) Sample PD9694l, BaSISS mutation signal segregation patterns echo those seen in PD9694d and support phylogenetic tree/genotype solution 1 (green and blue mutations co-localise: green and yellow mutations segregate; lower *FGFR1* expression in green mutation areas).

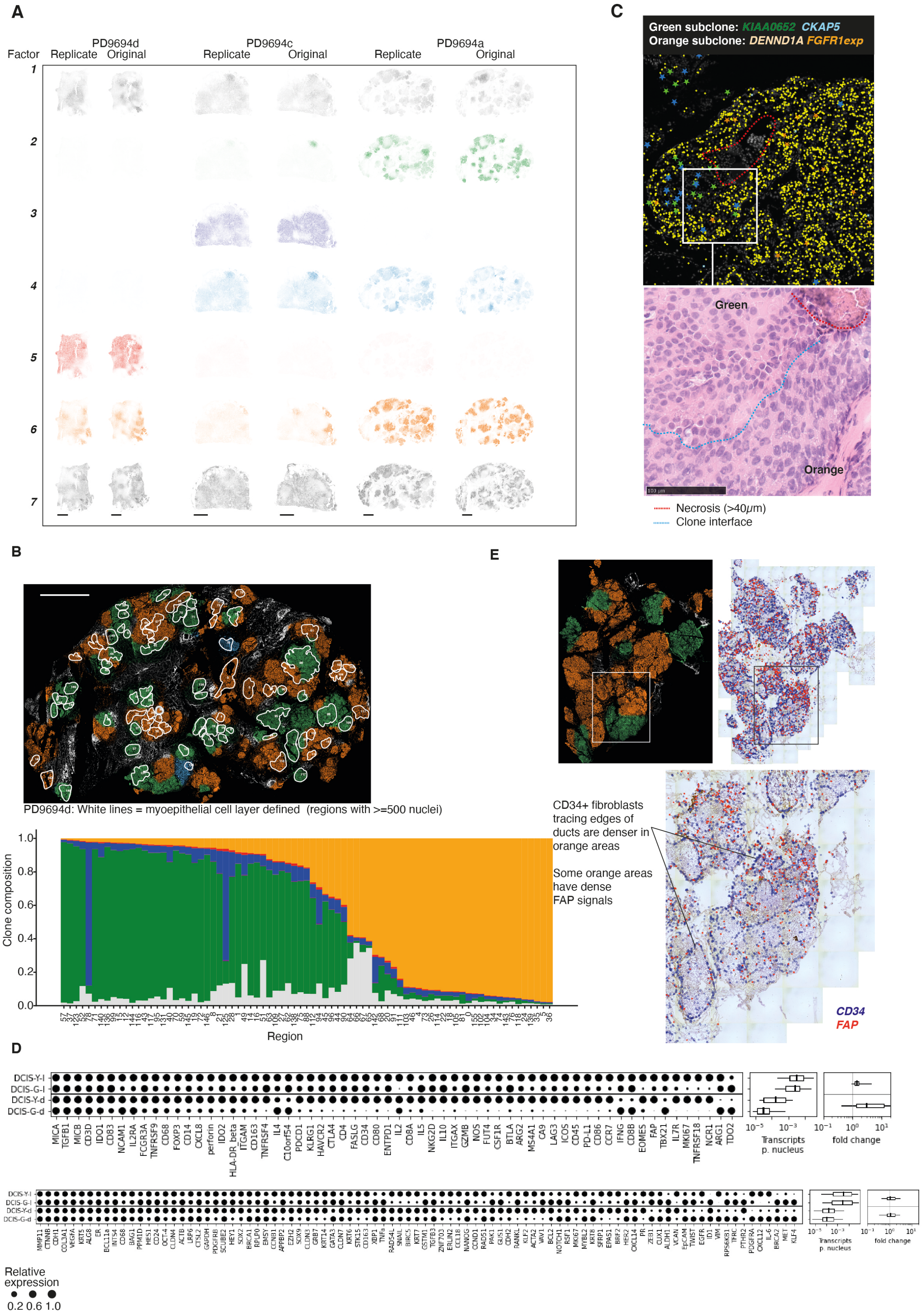

A
PD9694d
PD9694c
PD9694a
Factor
Replicate
Original
B
PD9694d: White lines = myoepithelial cell layer defined (regions with >=500 nuclei)
Clone composition
Region
C
Green subclone: KIAA0652 CKAP5
Orange subclone: DENND1A FGFR1exp
Green
Orange
Necrosis (>40μm)
Clone interface
D
DCIS-Y-l
DCIS-G-l
DCIS-Y-d
DCIS-G-d
Transcripts p. nucleus
fold change
Relative expression
0.2 0.6 1.0
E
CD34+ fibroblasts tracing edges of ducts are denser in orange areas
Some orange areas have dense FAP signals
CD34
FAP

**Fig. S3. Case PD9694 clone specific characteristics.** (**A**) Spatial genomics defined clone fields derived from replicate sequencing data: Factor 1-6 are clones corresponding to the same coloured phylogenetic tree branch. Factor 7 is residual. Relates to Fig 2E. (**B**) Stacked barplot of cancer clone composition of 'spaces' in PD9694d. Anatomically contiguous spaces (acini and ducts) are manually defined according to the presence of myoepithelial borders and/or intervening stroma (white borders represented on clone map). Regions containing >=500 cells are included in the analysis. (**C**) Example of a small green clone region adjacent to an area of necrosis surrounded by orange clone. Histological appearances corroborate the BaSISS signals. (**D**) Dot plots and boxplots reveal relative expression of ISS signals in Green (G) and Orange (Y) clones in samples PD9694d and PD9694l. Expression is normalised by sample. Dot area = (transcript/nucleus) divided by maximum value for each gene. Boxplots report transcripts/ nucleus (left) for genes shown and fold change between indicated comparisons (right). For all boxplots we report median, lower and upper quartiles (box) and 5-95 percentiles (whiskers). Expression per clone was defined on 162 - 3262 tiles, see Table S6 for details. (**E**) Clone map in PD9694l, a pure DCIS sample. Displayed alongside is the tissue wise and focus area of the ISS general fibroblast marker, *CD34* and fibroblast activation protein, *FAP* that both show spatial enrichment for the orange clone regions.

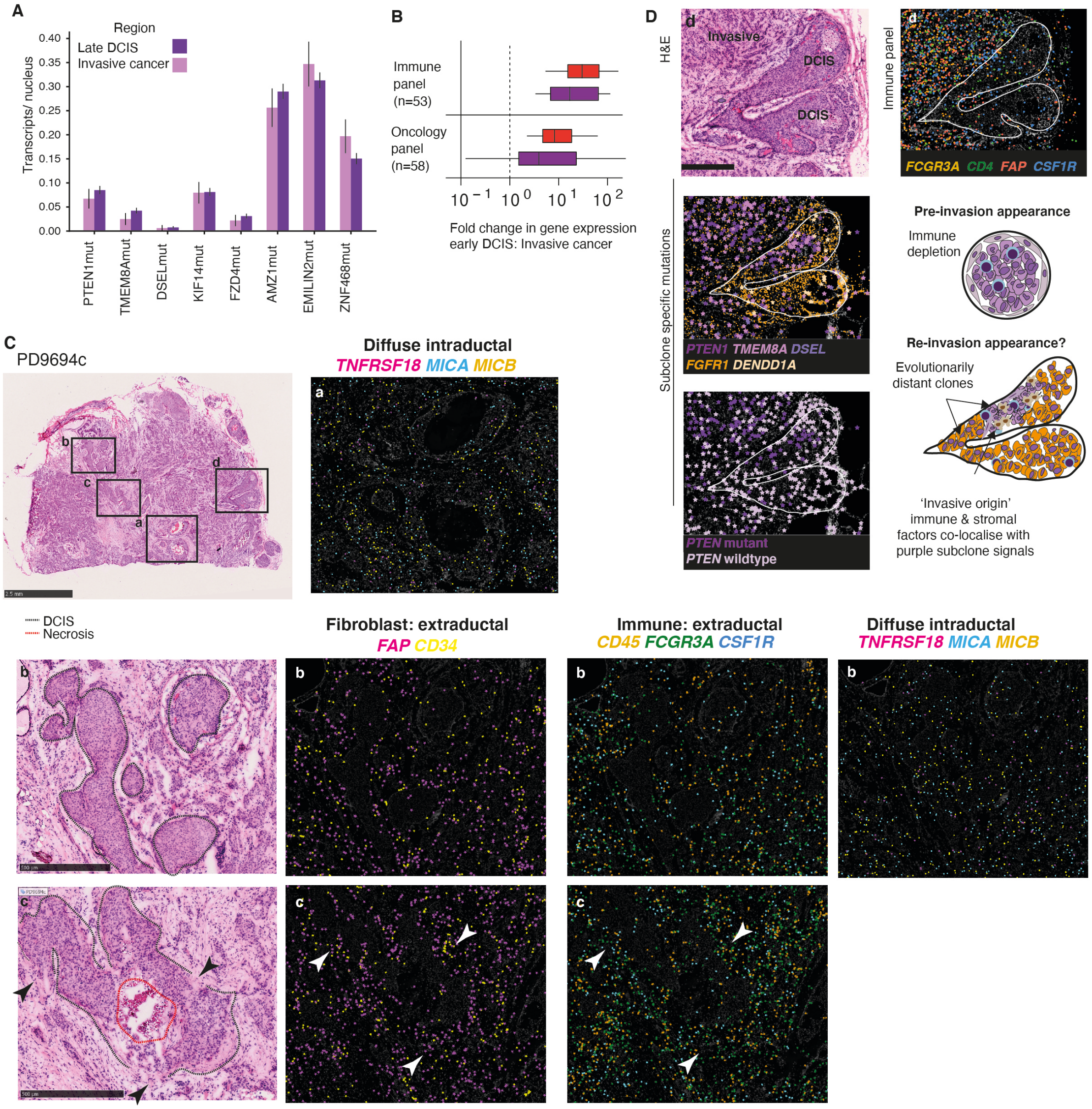

**Fig. S4. Temporal ordering during progression to invasion** (**A**) Barplots of purple subclone mutation signals in late DCIS and invasive cancer regions in PD9694c revealing similarity between the purple clone occupying both DCIS and invasive compartments (CI=5-95%). (**B**) Fold change in gene signals per nucleus in early DCIS (yellow or blue) and invasive cancer (red or purple) in the red and purple lineage respectively for the two ISS targeted gene panels. Significantly altered genes (PPLR<=0.01 with bonferroni adjustment) are included. (**C**) PD9694c H&E images (left) and select ISS expression signals (projected on DAPI background) to demonstrate differences in expression patterns between purple DCIS and purple invasive areas. In box a/b the myoepithelial border appears intact and most signals are mainly extraductal while some (far right) show a diffuse, similar distribution both inside and outside the ducts. In box c black/white arrowheads denote likely breaches in the ductal border and the associated ISS images show denser clusters of immune and fibroblast signals at this point probably 'entering' the duct. (**D**) Focus area of PD9694c (corresponds to S4C box d) region. This duct is predominantly occupied by the orange DCIS clone (Fig 3B-C) but a purple clone region is also present: left, middle and lower box reveal select purple subclone mutations (blue mutations are also present but not shown). Given the evolutionary distance between these clones (decades) this is surprising. The presence of clustering immune/ fibroblast signals (top right box) that colocalise specifically to the purple clone, this could be reinvasion of the duct by established invasive disease.

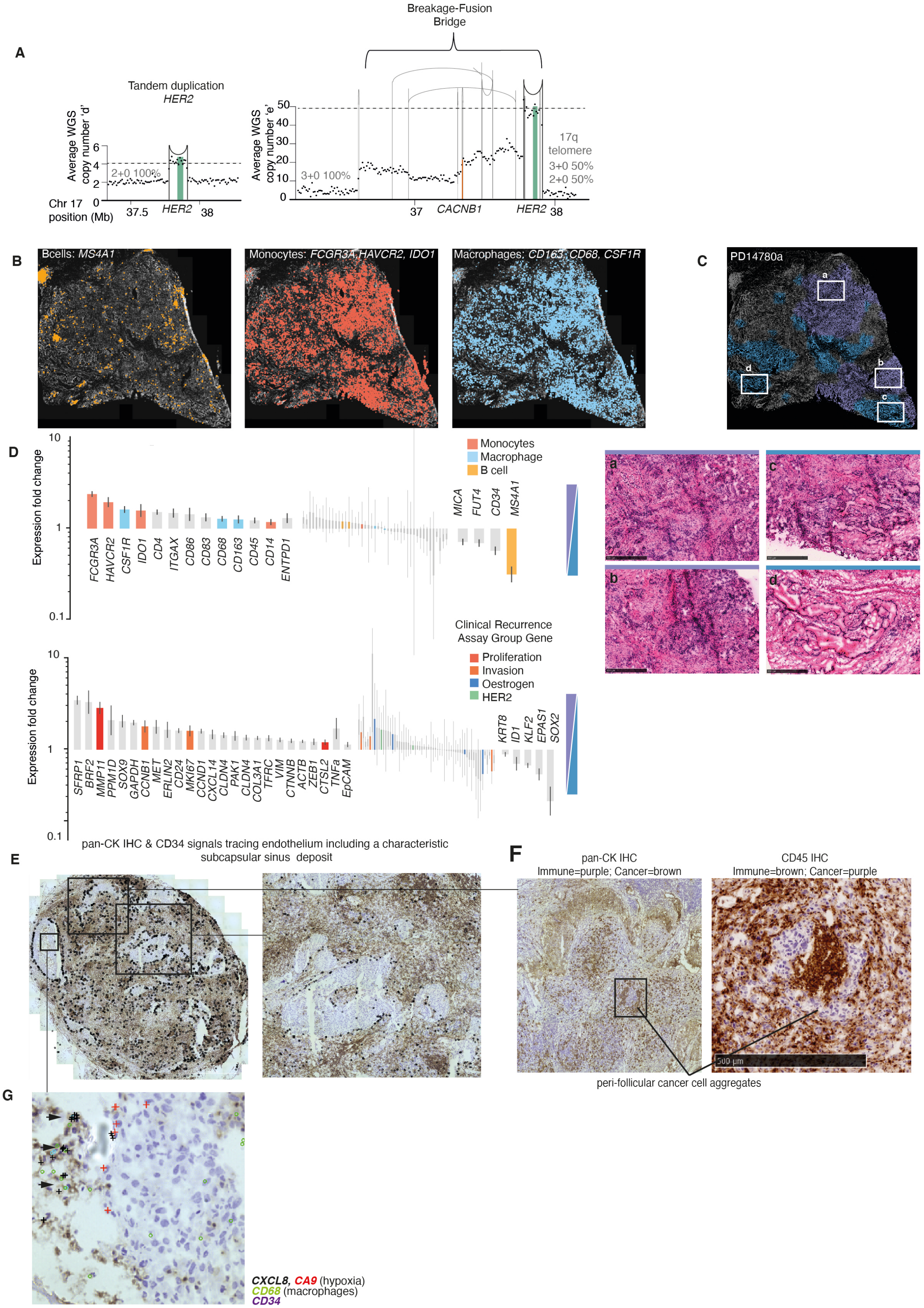
A
Tandem duplication
HER2
Average WGS copy number 'd'
2+0 100%
Chr 17 position (Mb)
37.5
HER2
38
Breakage-Fusion Bridge
Average WGS copy number 'e'
3+0 100%
17q telomere
3+0 50%
2+0 50%
37
CACNB1
HER2
38
B
Bcells: MS4A1
Monocytes: FCGR3A, HAVCR2, IDO1
Macrophages: CD163, CD68, CSF1R
C
PD14780a
D
Expression fold change
Monocytes
Macrophage
B cell
FCGR3A
HAVCR2
CSF1R
IDO1
CD4
ITGAX
CD86
CD83
CD68
CD163
CD45
CD14
ENTPD1
MICA
FUT4
CD34
MS4A1
Clinical Recurrence Assay Group Gene
Proliferation
Invasion
Oestrogen
HER2
SFRP1
BRF2
MMP11
PPM1D
SOX9
GAPDH
CCNB1
MET
ERLIN2
CD24
MKI67
CCND1
CXCL14
CLDN4
PAK1
CLDN4
COL3A1
TFRC
VIM
CTNNB
ACTB
ZEB1
CTSL2
TNFa
EpCAM
KRT8
ID1
KLF2
EPAS1
SOX2
E
pan-CK IHC & CD34 signals tracing endothelium including a characteristic subcapsular sinus deposit
F
pan-CK IHC
Immune=purple; Cancer=brown
CD45 IHC
Immune=brown; Cancer=purple
500 µm
peri-follicular cancer cell aggregates
G
CXCL8, CA9 (hypoxia)
CD68 (macrophages)
CD34

**Fig. S5. Spatial evolution in an invasive cancer and lymph node metastasis**

(**A**) Structural variant plots of *HER2* amplification. Vertical lines represent structural variant breakpoints. Dots represent local (binned) copy number. A tandem duplication (left plot) across the *HER2* locus occurs early in evolution and is shared by all samples. A mutation in the 3' UTR of one copy of the HER2 gene predates amplification. A breakage fusion bridge (BFB) amplifies the mutant and non-mutant version of HER2 and results in a novel CACNB1 fusion gene. (**B**) PD14780a ISS immune panel signals exhibit spatial variation and relates to the detected genetic clone distributions. (**C**) Histological appearances of blue and purple subclone regions are distinct. H&E images (fresh frozen tissue, 10um thick) reveal more duct formation in blue clone regions. (**D**) Barplot reporting expression fold change in ISS signals between the purple (838 tiles) and blue (838 tiles) clone. Significantly altered genes are shown (PPLR after bonferroni adjustment <=0.01). (**E**) PD14780e CD45 IHC image (immune cells = brown) with overlaid CD34 signals that trace the perimeter of many of the solid deposits. (**F**) Perifollicular growth pattern detected by pan-cytokeratin staining is confirmed by CD45 IHC – the cells are CD45- confirming that these are not ill-stained dendritic cells. (**G**) Focus image of hypoxia genes CA9 and CXCL8. CXCL8 colocalises CD45 IHC positive (brown stained) cells and macrophage markers CD68 or CD163.

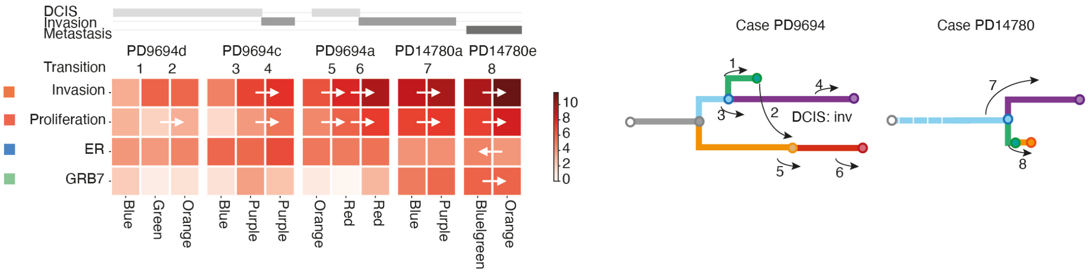

**Fig. S6. Heatmap of clinical recurrence assay gene group expression across 8 transition points in 2 cancers.** Each tile represents the relative expression of the gene ‘group’ as included in the 21 gene recurrence assay for breast cancer (Paik et al. *NEJM*, 2004): Invasion - *MMP11, CTSL2*; Proliferation = *CCNB1, MKI67, STK15, Survivin, MYBL2*; ER = *ER, PGR, BCL2, SCUBE2;* GRB7= *HER2, GRB7*. Relative weightings are described in Extended Methods section 8.5. Each ‘transition’ is represented by a number above the heatmap and corresponding phylogenetic tree and reports either a direct clonal succession event within a cancer lineage (1,3,5,7,8), the progression from DCIS (intraductal) to invasion (extraductal) (4,6) or competition from a divergent clone (2). White arrows indicate statistically significant fold changes between tiles (PPLR<0.01, Bonferroni adjusted). Proliferation changes consistently accompany the predicted pattern of clonal succession in 7 cases. The exception (transition 1) reflects the green pure DCIS clone that succeeded blue and it is likely that the blue clone has acquired additional unsampled mutations (below detection of WGS), furthermore, the green clone was clinically non-progressive.

#### Supplementary Tables

**Table S1.** (provided as Excel file)
**Padlock probe designs.**

**Table S2.** (provided as Excel file)
**Clinical and sample details.**

**Table S3.** (provided as Excel file)
**Bulk genomic sequence data summary.**

**Table S4.** (provided as Excel file)
**BaSISS target coverage.**

**Table S4.** (provided as Excel file)
**ISS by gene coverage.**

**Table S6.** (provided as Excel file)
**Clone specific expression comparisons.**
